# MATHEMATICAL MODELING OF RETINAL DEGENERATION: AEROBIC GLYCOLYSIS IN A SINGLE CONE

**DOI:** 10.1101/2020.07.06.190470

**Authors:** Erika Camacho, Atanaska Dobreva, Kamila Larripa, Anca Rǎdulescu, Deena Schmidt, Imelda Trejo

**Affiliations:** Arizona State University; North Carolina State University; Humboldt State University; SUNY New Paltz; University of Nevada, Reno; Los Alamos National Lab

**Keywords:** Retina, Photoreceptors, Aerobic Glycolysis, *β*-oxidation, Differential Equations

## Abstract

Cell degeneration, including that resulting in retinal diseases, is linked to metabolic issues. In the retina, photoreceptor degeneration can result from imbalance in lactate production and consumption as well as disturbances to pyruvate and glucose levels. To identify the key mechanisms in metabolism that may be culprits of this degeneration, we use a nonlinear system of differential equations to mathematically model the metabolic pathway of aerobic glycolysis in a healthy cone photoreceptor. This model allows us to analyze the levels of lactate, glucose, and pyruvate within a single cone cell. We perform numerical simulations, use available metabolic data to estimate parameters and fit the model to this data, and conduct a sensitivity analysis using two different methods (LHS/PRCC and eFAST) to identify pathways that have the largest impact on the system. Using bifurcation techniques, we find that the system has a bistable regime, biologically corresponding to a healthy versus a pathological state. The system exhibits a saddle node bifurcation and hysteresis. This work confirms the necessity for the external glucose concentration to sustain the cell even at low initial internal glucose levels. It also validates the role of *β*-oxidation of fatty acids which fuel oxidative phosphorylation under glucose- and lactate-depleted conditions, by showing that the rate of *β*-oxidation of ingested outer segment fatty acids in a healthy cone cell must be low. Model simulations reveal the modulating effect of external lactate in bringing the system to steady state; the bigger the difference between external lactate and initial internal lactate concentrations, the longer the system takes to achieve steady state. Parameter estimation for metabolic data demonstrates the importance of rerouting glucose and other intermediate metabolites to produce glycerol 3-phosphate (G3P), thus increasing lipid synthesis (a precursor to fatty acid production) to support their high growth rate. While a number of parameters are found to be significant by one or both of the methods for sensitivity analysis, the rate of *β*-oxidation of ingested outer segment fatty acids is shown to consistently play an important role in the concentration of glucose, G3P, and pyruvate, whereas the extracellular lactate level is shown to consistently play an important role in the concentration of lactate and acetyl coenzyme A. The ability of these mechanisms to affect key metabolites’ variability and levels (as revealed in our analyses) signifies the importance of inter-dependent and inter-connected feedback processes modulated by and affecting both the RPE’s and cone’s metabolism.

## 1 Introduction

Photoreceptors are the sensory cells of the eye, and they are the most energetically demanding cells in the body [52]. Photoreceptors have the most essential role in vision, absorbing light photons and processing them to electrical signals that can be transmitted to the brain. Therefore, vision deterioration or blindness occurs if the vitality and functionality of photoreceptors are compromised. In order to understand how to mitigate such pathological cases, it is essential to first obtain a firm grasp of processes that ensure the health of photoreceptors. The factor of upmost importance for photoreceptor vitality and functionality is metabolism.

To maintain their high metabolic demands and prevent accumulation of photo-oxidative product, the photoreceptors undergo constant renewal and periodic shedding of their fatty acid-rich outer segment (OS) discs. Aerobic glycolysis is integral to the renewal process. It facilitates the production of energy and the synthesis of phospholipids, both which are required for OS renewal. Phagocytosis of the shed OS by the retinal pigment epithelium (RPE) contributes to the creation of intermediate metabolites fundamental for photoreceptor energy production via *β*-oxidation [1]. Understanding the dynamics of glucose and lactate levels in aerobic glycolysis in a single cone cell is essential to maintain cone functionality and hence to preserve central vision. Studies in rod-less retinas have shown that maintaining functional cones even when 95% are gone may stop blindness [11, 30]. The purpose of this study is to analyze the key mechanisms affecting the levels of glucose, pyruvate, and lactate in a single cone cell via a first approximation mathematical model, with the goal of gaining insight into the interplay of glucose consumption and lactate production and consumption that may affect normal cone function.

### 1.1 Biological Background and Modeling Assumptions

#### 1.1.1 Photoreceptors and Retinal Pigment Epithelium (RPE)

Photoreceptors are specialized neurons that convert light into electrical signals that can be interpreted by the brain [37]. There are two types of photoreceptor cells: rods and cones. Cones are densely packed in the center of the retina and are responsible for color vision and high acuity. Rods have high sensitivity to light, are distributed on the outer edges of the retina, and are responsible for night and peripheral vision. In the human retina, there are approximately 90 million rod cells and 4.5 million cone cells [15] making rods twenty times more prevalent than cones. In the mature human retina (by about age 5 or 6), there are no spontaneous births of photoreceptors, making their preservation and vitality critical [10]. Photoreceptor shedding and renewal of their OS has been considered as a type of death and birth process as it is the mechanism by which photoreceptors discard unwanted elements (e.g., accumulated debris or toxic photo-oxidative compound in shed OS discs) and renew themselves through the recycling of various products. This process is a measurement of the photoreceptor’s energy uptake and consumption and associated metabolism [12, 11]. The shedding and renewal process and the associated metabolism of photoreceptors involve the RPE. The photoreceptors and the RPE work as a functional unit; glucose is transported from the RPE to the photoreceptors for their metabolism, lactate produced by photoreceptors and other retina cells is shuttled to the RPE for its metabolism, and the RPE mediates the phagocytosis of photoreceptor OS and recycling of fatty acids from these OS discs which are utilized in oxidative phosphorylation (OXPHOS) in the production of acetyl coenzyme A (ACoA); see Figure 1. However, as a first approximation we will not consider the role of the RPE but instead integrate the feedback mechanisms back into the cone cell via *β*-oxidation and external lactate transport.

**Figure 1:**
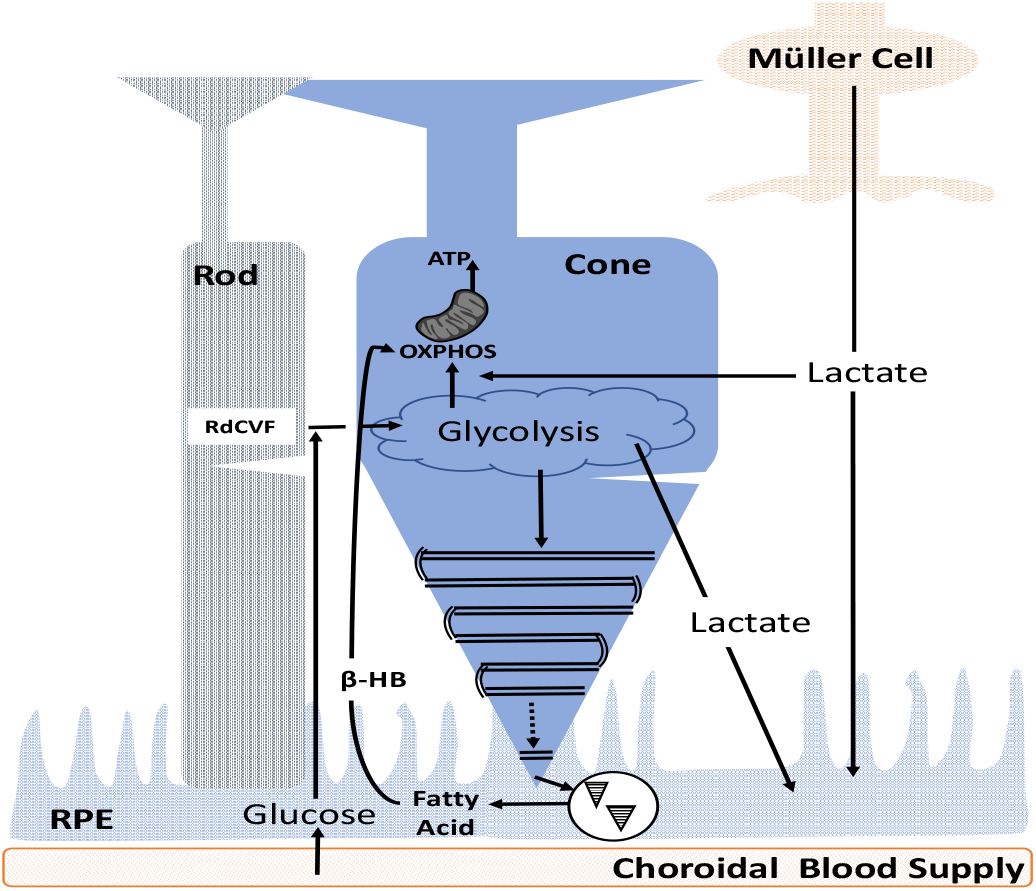
Schematic of metabolic pathways and substrate sources in the photoreceptors. This schematic shows that glucose and lactate flow between the cone photoreceptor and the RPE cell layer, as well as the role of glycolysis in providing energy to the cone cell and its role in helping generate cone outer segments. *β*-Hydroxybutyrate oxidation (*β*-HB) comes from oxidation of fatty acids from the shed outer segments so that under starvation or low glucose levels they can be used as oxidative substrates [1].

The RPE lies between the choroid and a layer of photoreceptors. In addition to functioning as the outer blood retinal barrier and transporting glucose to photoreceptor cells through GLUT1 (a facilitated glucose transporter), the RPE is involved in the phagocytosis of photoreceptor OS discs [57]. It serves as the principal pathway for the exchange of metabolites and ions between the choroidal blood supply and the retina [14]. Müeller cells are a layer of retinal glial cells and also provide support to photoreceptors. They can release lactate which is metabolized by photoreceptors [41] and store glycogen which can be broken down to glucose. A thorough investigation should consider the interaction of the three cell types. However in this work we consider, as a first step, a single cone photoreceptor in the human retina and model the metabolic pathways present. This analysis provides the foundation for a future application of the model: prediction of the interplay of metabolites from three cell types (RPE, photoreceptors, and Müeller) coexisting in the retina.

#### 1.1.2 Glycolysis and Oxidatative Phosphorylation

Photoreceptors are responsible for the majority of the energy consumption in the retina [38, 50]. Active transport of ions against their electrical and concentration gradients in neurons is required to repolarize the plasma membrane after depolarization, and this process is what consumes the most energy in photoreceptors [52, 37]. Moreover, the continual renewal and periodic shedding of OS [56] is also an extremely energetically demanding process.

All life on Earth relies on adenosine triphosphate (ATP) in energy transfer. ATP is produced via two pathways, oxidative phospholylation and glycolysis. Glycolysis, through a series of reactions (described in detail in Section 2.1), converts one molecule of glucose into two molecules of pyruvate, yielding two net molecules of ATP. If oxygen is present, pyruvate is typically converted to ACoA and enters the tricarboxylic acid (TCA) cycle, generating 32 net ATP molecules through OXPHOS. If oxygen is scarce, or if a cell has been metabolically reprogrammed, pyruvate is instead converted to lactate. However, photoreceptor cells use both pathways for energy production in the presence of oxygen with the vast majority of pyruvate being converted into lactate. In other words, despite only producing two molecules of ATP (versus 32 via OXPHOS), photoreceptors go through glycolysis as well as OXPHOS.

Glucose serves as the primary fuel in photoreceptors [13] and is broken down through aerobic glycolysis (glycolysis even in the presence of oxygen), termed the Warburg effect [2]. The Warburg effect has long been noted as a hallmark of tumors [23], but is also present in healthy tissue, particularly if their biosynthetic demands are high. Aerobic glycolysis maintains high fluxes through anabolic pathways and creates excess carbon which can be exploited for generation of nucleotides, lipids, and proteins, or diverted to other pathways branching from glycolysis, such as the pentose phosphate pathway and Kennedy pathway [32].

During glycolysis, glucose is transported into the cell. Rod-derived cone viability factor (RdCVF), which is secreted by rod photoreceptors, accelerates the uptake of glucose by cones through its binding with the glucose transporter complex 1/Basigin-1 (GLUT1/BSG-1) and stimulates aerobic glycolysis [3]. RdCVF also protects cones from degeneration [28], [53]. When glucose is in short supply, photoreceptors have the ability to take up and metabolize lactate [41].

#### 1.1.3 Lactate Secretion and Consumption

Photoreceptors can produce lactate from pyruvate and secrete it out of the cell or consume external lactate and convert it to pyruvate for OXPHOS if there is too much lactate in the extracellular space. The influx of lactate from the extracellular space would almost certainly slow the rate of glycolysis in the cell because any resulting higher intracellular lactate concentration shifts the lactate dehydrogenase (LDH)-catalyzed reaction equilibrium toward a higher NADH/NAD^+^ ratio. Under normal conditions, retinal cells oxidize cytosolic NADH to NAD^+^ (via the reduction/conversion of pyruvate to lactate in order to regenerate the NAD^+^). This lactate is transported out of the cell, thus increasing the amount of extracellular lactate. When glucose is low, such as during hypoglycemia or aglycemia conditions or hypoxia, oxidation of external lactate and fatty acids (via *β*-oxidation) to generate ACoA and thus produce energy (ATP) is favored [51]. When the photoreceptor cell undergoes OXPHOS, it makes citrate which provides an inhibitory feedback to glycolysis when other intermediates for ATP production are high, indicating additional glucose is not needed.

Glucose from the choroidal blood passes through the RPE to the retina where photoreceptors convert it to lactate, and in return, photoreceptors then export lactate as fuel for the RPE and for neighboring cells [26]. It has been hypothesized that photoreceptors also take up lactate for energy under low glucose levels. In humans, insufficient lactate transported out of the cone and rod cells for RPE consumption can suppress transport of glucose by the RPE. In such a case, the RPE takes glucose for its metabolism thereby decreasing the amount of glucose that is transported to the photoreceptors. Thus, lactate secretion for RPE consumption and external lactate consumption by photoreceptors is a balance process.

#### 1.1.4 Modeling Assumptions

Our model consolidates some of the steps in the glycolytic pathway in a single cone, for simplicity. Glucose is initially transported into the cell, and the rate of transport is amplified by the release of RdCVF from rods. The rate of transport is gradient dependent and modulated by the difference in the amount of glucose inside and outside the cell. The next step is the conversion of glucose in the cell into glucose-6-phosphate (G6P) by the enzyme hexokinase 2. This phosphorylation also works to trap glucose in the cell’s cytosol. Some G6P is diverted to the pentose phosphate pathway (not included in our model), while the rest moves through the glycolytic pathway. The enzyme phosphofructokinase (PFK) converts fructose-6-phosphate to fructose 1,6-biphosphate (not explicitly included in our model). This in turn is cleaved into two sugar molecules, one of which is dihydroxyacetone phosphate (DHAP), the substrate for the next reaction. DHAP is converted to glycerol-3-phosphate (G3P) in the Kennedy pathway and glyceraldehyde-3-phosphate (GAP) in the glycolytic pathway. The latter metabolite is not explicitly considered in our model. A number of sequential reactions occur, with the ultimate step aided by the enzyme pyruvate kinase, resulting in pyruvate. Since our model considers a single cone, we use the presence of the metabolite concentration [G3P] with an appropriate scaling factor as a proxy for the amount of RdCVF synthesized by the rods. RdCVF accelerates glucose uptake in cones [3, 28, 53].

Specifically, our model incorporates the uptake and consumption of glucose, the production of G3P and pyruvate, and key consecutive chemical reactions in the cone cell involving lactate, ACoA, and citrate. Pyruvate is converted to lactate, which is then transported out of the cell. A portion of pyruvate is also transferred to the mitochondria; there it is converted to ACoA and goes through OXPHOS, creating citrate, which leads to the production of ATP but also negatively regulates the glycotic pathway. Citrate inhibits phosphofructokinase (PFK) which slows down the production of G3P and pyruvate. G3P leads to the production of lipids which are used to create OS that are shed and phagocytized periodically. The fatty acids from the shed OS can feedback into the cone cell as *β*-hydroxybutyrate (*β*-HB) which serves as a substrate for ACoA production. Through the process of *β*-oxidation fatty acids in the RPE result in *β*-HB. The specific pathways are outlined in Figure 2, and specific evidence is presented for each pathway in detail below.

**Figure 2:**
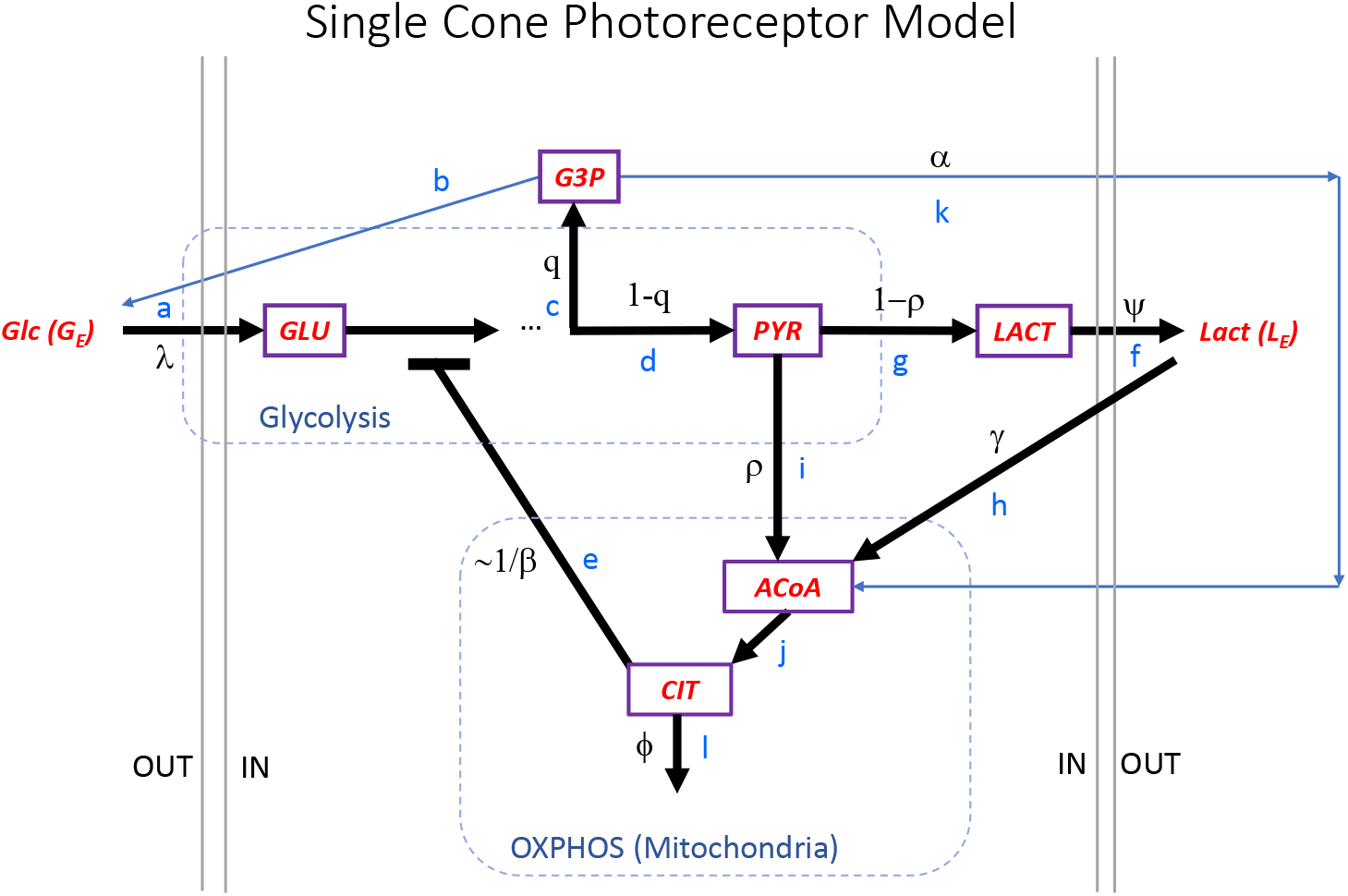
Flow diagram of the key metabolic pathways within a single cone photoreceptor. Parameters corresponding to each pathway are labeled with black letters while metabolic pathways are labeled with blue letters (a-l). Parameters are described in Table 2. Brief descriptions of each pathway are given in Table 1 and are described in detail in Section 2.1.

## 2 Mathematical Model

We model six key steps in the glycolytic pathway as a system of six nonlinear ordinary differential equations that describes metabolic pathways in a single cone. Specifically, we track the temporal dynamics of the following six concentrations in the cell: internal glucose ([G]), glyceraldehyde 3-phosphate ([G3P]), pyruvate ([PYR]), lactate ([LACT]), acetyl coenzyme A ([ACoA]), and citrate ([CIT]). The chemical reactions and up and down-regulations included in this model are illustrated in Figure 2 and listed in Table 1. In Section 2.1, we discuss the biological basis for each interaction pathway used in the model. In Sections 2.2 and 2.3, we give the model equations and parameter values used, respectively.

**Table 1:** Description of metabolic pathways in the model.

| Pathway | Description | References |
| --- | --- | --- |
| **a** | gradient transport of glucose | [40, 33] |
| **b** | glucose uptake; without and with RdCVF | [53, 3, 10] |
| **c** | glycolytic flow diverted to G3P | [43, 46] |
| **d** | glycolytic flow diverted to pyruvate | [6] |
| **e** | glycolysis inhibition by citrate | [6] |
| **f** | gradient gating mechanism to transport lactate out of the cell | [5, 9, 25, 8, 22] |
| **g** | fraction of pyruvate concentration converted into lactate | [18, 41, 55] |
| **h** | gradient gating mechanism to transport lactate into the cell for ACoA production | [20, 6] |
| **i** | fraction of pyruvate concentration converted into ACoA | [47, 36] |
| **j** | conversion of ACoA to citrate | [47] |
| **k** | β-HB utilized in production of ACoA | [1] |
| **l** | citrate is diverted to the cytosol and other metabolic pathways | [47] |

### 2.1 Kinetic pathways in the model

Here, we provide details of all model pathways shown in Figure 2 and described in Table 1. These pathways represent a reduced system, with some pathways omitted and elements implicitly modeled via proxies. There are multiple intermediates produced in glycolysis and oxidative phosphorylation which are not explicitly considered in this work. In order to focus on production and consumption of glucose, lactate, and pyruvate in a single cone cell we reduce the system to its most essential components and pathways.

#### Pathway a: gradient transport of glucose

In the retina, sodium independent glucose transporters (GLUTs) transport glucose by facilitated diffusion down its concentration gradient [40]. GLUT1 is found in human photoreceptor outer segments [33]. We model this pathway by considering the difference between external glucose concentration (the parameter *G_E_* in our model) and the internal glucose concentration, the variable [G]. The parameter λ is a constant of proportionality that governs the rate of glucose uptake based on the concentration gradient.

#### Pathway b: glucose uptake without and with stimulation of GLUT1 by RdCVF

Rod-derived cone viability factor (RdCVF) is secreted in a paracrine manner by rod photoreceptors and protects cones from degeneration [3], [53]. It binds with the GLUT1/BSG-1 complex to activate GLUT1 and accelerates the entry of glucose into the cone. We use [G3P] together with an appropriate scaling factor incorporated into *δ* as a proxy for the RdCVF that is synthesized by the rods. G3P is needed for phospholipid synthesis resulting in the renewal of photoreceptor OS [10]. Since for every cone cell there are approximately 20 rods in the human retina and G3P in our model is a measurement of the cone OS, we scale concentrations to account for rods’ secretion of RdCVF that accelerates glucose uptake and supports cone vitality.

Note that the parameter *n* is included in this term so that even in the absence of RdCVF or our proxy for rods, glucose is passively transported down its gradient (bidirectionally) into the intracellular space of the cone cell. Thus, λ*n* is the glucose uptake rate of our cone cell in the absence of RdCVF. The uptake of glucose due to RdCVF is an allosteric reaction, and therefore there is a binding time requirement for the enzyme to catalyze the reaction. We therefore use a Hill type function with a Hill coefficient of 2 to model the sigmoidal response [10], [42].

#### Pathway c: glycolytic flow diverted to G3P

The sequence of reactions leading to G3P are glucose to glucose-6 phosphate to fructose 6-phosphate to fructose 1,6-biphosphate to dihydroxyacetone phosphate (DHAP), and then G3P. Figure 2 indicates the many intermediate steps which are skipped with the ellipsis in the diagram. We model the conversion of glucose to G3P with a Hill type function where *V*_[G3P]_ is the maximal rate of conversion (controlled by the rate-limiting allosteric enzyme PFK described above) of glucose to G3P and *K*_[G3P]_ is the concentration of the ligand that gives half-maximal activity [46].

#### Pathway d: glycolytic flow diverted to pyruvate

We skip intermediate reactions to focus on the key metabolites of interest; glucose, pyruvate, and lactate. We take a similar approach as in pathway **c** and consider glucose to be the substrate in the reaction resulting in the production of pyruvate. We can infer from known aerobic glycolysis that the substrate (in this case glucose) which is not converted to G3P is converted to pyruvate [6]. Thus, a fraction *q* of glucose gets converted to G3P, while the remaining fraction 1 − *q* gets converted to pyruvate.

#### Pathway e: glycolysis inhibition by citrate

The flux through the glycolytic pathway must be responsive to conditions both inside and outside the cell, and the enzyme phosphofructokinase (PFK) is a key element in this control. PFK is inhibited by citrate, which enhances the allosteric inhibitory effect of ATP [6]. Elevated citrate levels indicate that biosynthetic precursors are readily available and additional glucose should not be degraded. The form of the function capturing this inhibition is reciprocal to the concentration of citrate and is multiplied to the metabolic reactions that involve glucose as a substrate down the glycolysis pathway (i.e., the reactions that produce G3P and pyruvate).

#### Pathway f: gradient gating mechanism to transport lactate out of the cell

Monocarboxylate transport proteins (MCT) are a family of plasma membrane transporters and allow lactate, pyruvate, and ketone bodies to be actively transported across cell membranes [5]. The RPE expresses various isoforms of the MCT transporter [9], as do the photoreceptors, Müeller cells, and the inner blood-retinal barrier. Inhibition of MCT results in retinal function loss [9], mainly due to lactate accumulation in the extracellular space. The lactate transport rate is dependent on pH, temperature, and concentration of internal lactate relative to external cellular lactate [25]. MCTs faciliate lactate transport down the concentration and pH gradients [8]. MCT1 is particularly important for reducing conditions of intracellular acidification when glycotic flux is high [22]. MCT1 transports lactate out of the photoreceptors and into the RPE. We incorporate this process in our model with pathway **f**.

Lactate flux out of the cell depends on the concentration gradient, pH, and temperature. We model this using a gating function *f* ([LACT]); see Equation 8. For large binding affinity of lactate transporter (i.e., for large *k* values), if the external lactate concentration *L_E_* exceeds the internal lactate concentration [LACT], the gate closes, and the external lactate is directed to OXPHOS via function *h*([LACT]) to produce ACoA; see Equation 9. The height of this function represents the maximal flux possible, physiologically limited by the concentration and expression of MCT1.

#### Pathway g: fraction of pyruvate concentration converted into lactate

Pyruvate is converted to lactate in glycolysis. This metabolic reaction is promoted by increased expression of the enzyme lactate dehydrogenase A (LDHA) and inactivation of pyruvate dehydrogenase [18], [41]. The conversion and direction of the reaction from pyruvate to lactate depends on lactate dehydrogenase subtypes; photoreceptors express LDHA which favors the production of lactate from pyruvate [10].

#### Pathway h: gradient gating mechanism to transport lactate into the cell for ACoA production

Pyruvate dehydrogenase complex (PDC) converts pyruvate to ACoA. The consumption of lactate back into the cell depends on a gating mechanism modulated by the pH levels and the lactate gradient inside and outside the cell. While LDHA converts pyruvate to lactate, lactate dehydrogenase B (LDHB) converts lactate to pyruvate. The latter reaction involves external lactate and the newly acquired pyruvate does not convert back to lactate but rather goes into the mitochondria where it becomes a substrate in the production of ACoA. The conversion of lactate to pyruvate and vice versa also depends on NAD^+^ and NADH levels as they can drive things in one direction or another. When lactate is used as an energy source, lactate carbon is ultimately inserted into the TCA cycle in the mitochondria.

Glycolysis and gluconeogenesis are coordinated so that within one cell, one pathway is relatively inactive while the other is highly active. The rate of glycolysis is governed by the concentration of glucose whereas the rate of gluconeogenesis is governed by the concentration of lactate [20]. Inhibition of the enzyme PFK (which drives glycolysis) and abundance in [CIT] activates gluconeogenesis [6]. Rather than modeling all steps of gluconeogenesis, we let external lactate feed directly to ACoA and do not track its passage through pyruvate. This mechanism consolidates entry of lactate into the cell.

#### Pathway i: fraction of pyruvate converted into ACoA

After pyruvate is produced, its flux branches off and a fraction *ρ* of pyruvate is transferred to the mitochondria by the mitochondrial pyruvate carrier and converted into ACoA. During glycolysis, the mitochondrial pyruvate dehydrogenase complex catalyzes the oxidative decarboxylation of pyruvate to produce ACoA [36], [47].

#### Pathway j: conversion of ACoA to citrate

In the mitochondria, the enzyme citrate synthase catalyzes the conversion of ACoA and oxaloacetate into citrate [47].

#### Pathway k: fatty acids utilized in production of ACoA

During the shedding and subsequent phagocytosis of the OS, a source of fatty acids is created [1]. This itself can be used for metabolism, and feeds directly into ACoA. As depicted in pathway **b**, we use G3P as a proxy for the substrates that are created through *β*-oxidation (the process by which fatty acid molecules are broken down in the mitochondria to generate ACoA). G3P is converted to lipids which form the photoreceptor’s OS that eventually get shed and become phagolysosomes containing fatty acids. These fatty acids can be oxidized and generate ACoA. ACoA leads to the production of *β*-Hydroxybutyrate (*β*-HB) via ketogenesis which can be used as an oxidative substrate in the TCA cycle when glucose is low. In conditions of glucose starvation, fatty acids are released, broken down, oxidized, and used to produce ketones that can be used to fuel the cone cell. In our mathematical model, we do not directly model ketogenesis but instead G3P serves as a proxy for *β*-oxidation of fatty acids from ingested OS.

#### Pathway l: citrate is diverted to the cytosol and other metabolic pathways

Citrate in the mitochondria can be oxidized via the TCA cycle, or it can be moved to the cytosol to be cleaved by ATP citrate lyase, which regenerates ACoA and oxaloacetate. This pathway redirects ACoA away from the mitchondria under conditions of glucose excess [47]. It reduces the glycolytic flux coming into the TCA cycle and signals the cone cell that ATP is high and there is no need for glucose metabolism.

### 2.2 Model Equations

Following the flow diagram given in Figure 2, we apply mass-action Michaelis-Menten kinetics and allosteric regulations to the relevant parts of the variable interactions to yield the resulting system of equations:

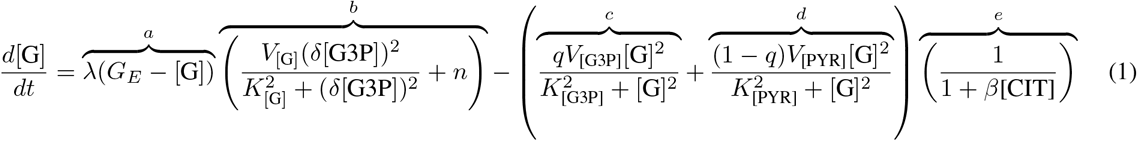

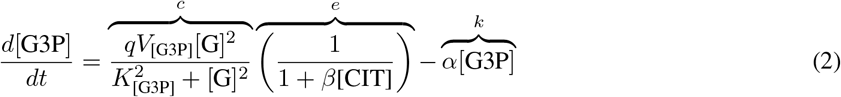

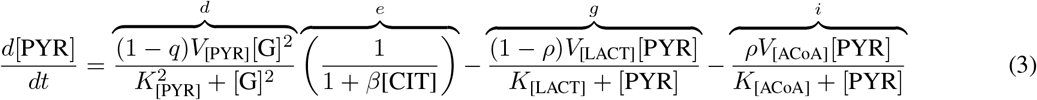

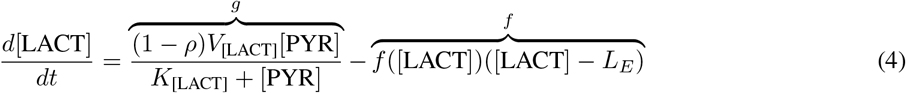

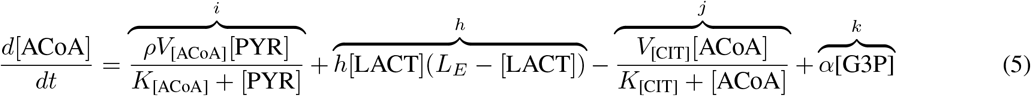

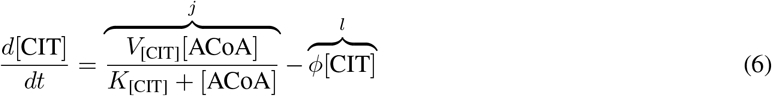

The model consists of 25 parameters defining various metabolic kinetic processes affecting internal [G], [PYR], and internal [LACT] within a cone cell; see Table 2. Since we are not incorporating the RPE and the rod cells, we consider three intermediate metabolites, G3P, ACoA, and citrate, that affect energy production and are sources of feedback mechanisms. The former two provide feedback mechanisms for glucose and fatty acids (in the form of *β*-HB) to enter the cone cell. They are proxies for mechanisms being mediated by the RPE and rod cells. The metabolite G3P in a healthy cone cell can be used to approximate the rods that synthesize RdCVF as well as the fatty acids that are *β*-oxidized, converted to *β*-HB, and contribute to ACoA. The intermediate metabolite ACoA is a product of pyruvate and OS fatty acids and is the entry point of the citric acid cycle, also known as the Krebs cycle or tricarboxylic acid (TCA) cycle. Citrate provides a self-regulating mechanism through its inhibition of PKF. If citrate builds up, it signals the cell that the citric acid cycle is backed up and does not need more intermediates to create ATP, slowing down glycolysis. This in turn reduces the production of pyruvate and lactate. The six key metabolic processes under consideration in this study are described by equations (1)-(6) and the 25 parameters, following key features of photoreceptor biochemistry [10, 31, 29]. As such we define glycerol-3-phosphate as G3P, which should not be confused with glyceraldehyde-3-phosphate (abbreviated as GAP, G3P, and GA3P in some literature).

**Table 2:** Parameter descriptions and units.

| Parameter | Description | Units |
| --- | --- | --- |
| $\lambda$ | Transport conversion factor | $\text{mM}^{-1}$ |
| $V_{[\text{G}]}$ | Maximum transport rate of glucose | $\text{mM} \cdot \text{min}^{-1}$ |
| $K_{[\text{G}]}$ | Substrate concentration that gives half the maximal rate of $V_{[\text{G}]}$ | $\text{mM}$ |
| $n$ | Rate of passive glucose transport in the absence of RdCVF | $\text{mM} \cdot \text{min}^{-1}$ |
| $\delta$ | Scaling factor for contribution of RdCVF by rods | no units |
| $q$ | Fraction of G converted into G3P | no units |
| $V_{[\text{G3P}]}$ | Maximum production rate of G3P | $\text{mM} \cdot \text{min}^{-1}$ |
| $K_{[\text{G3P}]}$ | Substrate concentration that gives half the maximal rate of $V_{[\text{G3P}]}$ | $\text{mM}$ |
| $V_{[\text{PYR}]}$ | Maximum production rate of PYR | $\text{mM} \cdot \text{min}^{-1}$ |
| $K_{[\text{PYR}]}$ | Substrate concentration that gives half the maximal rate of $V_{[\text{PYR}]}$ | $\text{mM}$ |
| $\beta$ | Rate of CIT inhibition of G catabolism (times an appropriate conversion factor) | $\text{mM}^{-1}$ |
| $\alpha$ | Rate of β-oxidation of ingested OS fatty acids (created from G3P) to generate the β-HB substrate for ACoA | $\text{min}^{-1}$ |
| $\rho$ | Fraction of PYR converted into ACoA | no units |
| $V_{[\text{LACT}]}$ | Maximum production rate of LACT | $\text{mM} \cdot \text{min}^{-1}$ |
| $K_{[\text{LACT}]}$ | Substrate concentration that gives half the maximal rate of $V_{[\text{LACT}]}$ | $\text{mM}$ |
| $V_{[\text{ACoA}]}$ | Maximum production rate of ACoA | $\text{mM} \cdot \text{min}^{-1}$ |
| $K_{[\text{ACoA}]}$ | Substrate concentration that gives half the maximal rate of $V_{[\text{ACoA}]}$ | $\text{mM}$ |
| $\phi$ | Rate of CIT converted to ATP | $\text{min}^{-1}$ |
| $V_{[\text{CIT}]}$ | Maximum production rate of CIT | $\text{mM} \cdot \text{min}^{-1}$ |
| $K_{[\text{CIT}]}$ | Substrate concentration that gives half the maximal rate of $V_{[\text{CIT}]}$ | $\text{mM}$ |
| $\psi$ | Maximum velocity of lactate transport | $\text{min}^{-1}$ |
| $k$ | Measurement of binding affinity of lactate transporter | $\text{mM}^{-1}$ |
| $\gamma$ | Maximum velocity of lactate transport contributing to ACoA | $\text{min}^{-1}$ |
| $G_E$ | Concentration of glucose outside the cell | $\text{mM}$ |
| $L_E$ | Concentration of lactate outside the cell | $\text{mM}$ |

Equation (1) describes the rate of change with respect to time of the glucose concentration. It increases or decreases proportional to bidirectional glucose transport and decreases by catalysis. The transport function of [G3P] [10]:

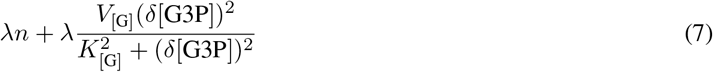

accounts for the passive transport term (first term of Equation (7)) and the facilitated transport term (second term of Equation (7)). In passive transport, glucose crosses the membrane without activation and stimulation by the facilitated transporter GLUT1, while in facilitated transport, RdCVF stimulates the transport activity of GLUT1 by triggering its tetramerization and accelerating the uptake of glucose [10]. The expression *δ*[G3P] accounts for the concentration of RdCVF synthesized by rod phothoreceptors since it is assumed that RdCVF concentration is in proportion to [G3P].^1^

We model *q* as the fraction of [G] that is converted into [G3P] and 1 − *q* as the remaining fraction of [G] that is converted into [PYR]. The metabolism of glucose into these two metabolites is inhibited by [CIT], where *β* is the rate of citrate inhibition of glucose catabolism.

Equation (2) describes the rate of change with respect to time of the G3P concentration. [G3P] increases with an influx of glucose, which is inhibited by citrate, and decreases by production of OS, which serves as a measurement of *β*-oxidation of ingested OS fatty acids that contribute to the production of ACoA. We are taking catabolism of *α*[G3P] as a proxy for OS fatty acids converted into ACoA.^2^

Equation (3) describes the rate of change with respect to time of the pyruvate concentration. [PYR] increases with an influx of glucose, which is inhibited by citrate, and decreases by its conversion into lactate and ACoA. The factor (1 − *ρ*) accounts for the fraction of [PYR] converted into lactate while *ρ* accounts for the fraction of [PYR] converted into ACoA.

Equation (4) describes the rate of change with respect to time of the lactate concentration. [LACT] increases by conversion of pyruvate to lactate via aerobic glycolosis and increases or decreases by bidirectional lactate transport. The lactate transport rate is modeled with a logistic function as follows:

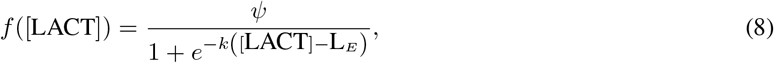

where *ψ* is the maximum transport rate, L_*E*_ is the extracellular concentration of lactate, and *k* is the binding affinity of lactate transporters which corresponds to the steepness of the curve *f*. Since L_*E*_ accounts for the lactate concentration outside of the cell, the gradient flux of lactate is from inside to outside of the cell when [LACT] > L_*E*_ while the opposite gradient flow occurs when [LACT] < L_*E*_. If external lactate is in abundance, then the transport rate out of the cell is very small, i.e.,

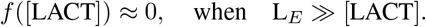

In other words, if lactate inside of the cell is scarce relative to external lactate then the transport of lactate out of the cell is a slow process. If the intracellular lactate concentration is much larger than the extracellular concentration, then lactate transport out of the cell is faster, i.e.,

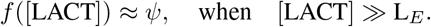

Equation (5) describes the rate of change with respect to time of the ACoA concentration. [ACoA] increases by PYR leakage to the mitochondria and *β*-HB produced from OS fatty acids generated by G3P lipid synthesis. It also increases or decreases by bidirectional lactate transport and decreases by its conversion into citrate. The lactate transport rate is modeled with a logistic function as follows:

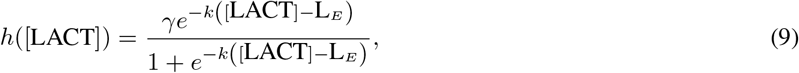

where *γ* is the maximum transport rate and *k* is the steepness of the curve *h*([LACT]).^3^ The extracellular lactate that comes into the cell gets converted into pyruvate which is immediately shuttled to the mitochondria for OXPHOS, and there is no re-conversion of lactate. Mathematically, this means that we can directly model the gradient influx of external lactate into the mitochondria and the conversion of this lactate to ACoA with the transport rate *h*([LACT]). The conversion of L_*E*_ to ACoA is metabolically faster when external lactate is in abundance, i.e.,

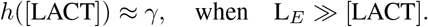

However, when the L_*E*_ is scarce, its contribution to the production of ACoA is negligible, i.e.,

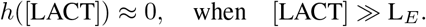

Equation (6) describes the rate of change with respect to time of the citrate concentration. [CIT] increases by the conversion of ACoA into citrate and decreases by its conversion into other intermediate metabolites leading to the creation of ATP.

In our model every resulting product becomes the substrate in the next metabolic reaction, with the exception of citrate, the last metabolite in our sequence of metabolic reactions, and lactate which is modulated by L_*E*_. The metabolic conversion of the substrates [G], [G3P], [PYR], and [ACoA] into their respective products, given by the variables in equations (1)-(6), are modeled with Hill type functions:

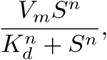

where *V_m_* is the maximal velocity of the reaction, *K_d_* is the dissociation constant (or equivalently concentration of the substrate at which the conversion rate achieves its half-maximum value) and *n* = 1, 2 is the Hill coefficient. This coefficient relates to the number of binding sites available in the enzyme. When there is cooperative binding, *n* is greater than one, illustrating higher binding affinity of the substrate to the enzyme [46]. We modeled allosteric regulation kinetics with *n* = 2 indicative of multiple binding sights and enzyme cooperation, which results in increased substrate conversion rates after the first binding event.

### 2.3 Parameter Values

All model parameters and their meaning are described in Table 2. We performed an extensive literature search to identify and justify parameter values and ranges used in the model; see Table 3. When human values were not available, we used animal values. Note that even through *V*_[G3P]_ and *V*_[PYR]_ have the same baseline values, their corresponding range values, used later for the sensitivity analysis, are different. When metabolic parameter values for retina cells were not available, we used values from brain, heart, liver, or muscle tissues. Cancer cells can also serve as a case study to investigate the predictive capabilities of our model, as they also exhibit the Warburg effect, converting glucose to lactate even in the presence of oxygen. Since both cancer and photoreceptor cells utilize aerobic glycolysis for metabolism and both are high energy demanding cells, we used cancer data to see how well our cone cell model extends to other aerobic glycolysis systems.

**Table 3:** Parameter values used in simulations for photoreceptor model and cancer cells.

| Parameter | Normal photoreceptor model |  | Cancer cells |  | Reference |
| --- | --- | --- | --- | --- | --- |
|  | Range Value | Baseline Value | Range Value | Baseline Value |  |
| $\lambda$ | [0.062, 0.093] | 0.0755 | [0.054, 0.101] | | [10]<sup>+</sup> & Est |
| $V_{[G]}$ | [0.1, 1.56] | 1.2 | | | [10, 54] |
| $K_{[G]}$ | [5, 24.7] | 19 | | | [10, 54]<sup>+</sup> |
| $n$ | [0.0007, 0.0013] | 0.001 | | | [10]<sup>+</sup> |
| $\delta$ | [45.5, 95] | 65 | | | [10]<sup>+</sup> |
| $q$ | [0.04, 0.2] | 0.18 | [0.336, 0.624] | 0.38 | [39]<sup>+</sup> & Est |
| $V_{[G3P]}$ | [0.12, 0.18] | 0.15 | | | [10] |
| $K_{[G3P]}$ | [0.02, 0.171] | 0.143 | | | [10]<sup>+</sup> |
| $V_{[PYR]}$ | [0.0013, 0.3915] | 0.15 | | | [10, 54] |
| $K_{[PYR]}$ | [0.05, 2.21] | 1.7 | | | [10, 54]<sup>+</sup> |
| $\beta$ | [0.7, 1.3] | 1 | [0.56, 1.04] | 0.8 | [19]<sup>+</sup> & Est |
| $\alpha$ | [0.002, 1] | 0.2 | | | [10]<sup>+</sup> |
| $\rho$ | [0.04, 0.06] | 0.05 | | | [10, 51, 21, 16, 17] |
| $V_{[LACT]}$ | [0.098, 0.33] | 0.14 | | | [10, 54] |
| $K_{[LACT]}$ | [0.0875, 10] | 0.125 | | | [10, 54]<sup>+</sup> |
| $V_{[ACoA]}$ | [0.105, 0.195] | 0.15 | | | [54] |
| $K_{[ACoA]}$ | [0.005, 0.02] | 0.02 | | | [35] |
| $\phi$ | [0.7, 1.3] | 1 | [0.35, 0.65] | 0.5 | [4] & Est |
| $V_{[CIT]}$ | [0.021, 0.039] | 0.03 | [0.07, 0.13] | 0.1 | [35, 4] & Est |
| $K_{[CIT]}$ | [0.0046, 0.00702] | 0.0054 | [0.35, 0.65] | 0.5 | [35] & Est |
| $\psi$ | [6.4, 9.6] | 8 | | | [10] |
| $k$ | [7, 13] | 10 | [0.315, 0.585] | 0.45 | CS & Est |
| $\gamma$ | [0.7, 1.3] | 1 | [0.0105, 0.0195] | 0.015 | [27] & Est |
| $G_E$ | [5, 20] | 11.5 | [8.4, 15.6] | 13 | [10, 54, 51] & Est |
| $L_E$ | [5, 22] | 10 | [15.4, 28.6] | 22 | [10, 54] & Est |

## 3 Numerical Results

### 3.1 Model Validation

With parameter values in empirical ranges, we first verified that the model predicts a temporal evolution comparable to that observed in data. To do this, we compared model simulations with results from an empirical study in cancer cells, which provided measurements of the intracellular concentrations of glucose, lactate, and pyruvate over a period of four hours [54]. Ying et al. [54] measured these concentrations at six time points (0 hours, 0.5 hours, 1 hours, 2 hours, 3 hours, and 4 hours). 4T1 (breast cancer line) cells were cultured in 10 mM of both glucose and lactate with a pH of 7.4. The concentrations of glucose, lactate, and pyruvate were measured using a spectrophotometer.^4^ We averaged the experimental results and used the resulting data, given in Table 4, as a first step in validating our model for an aerobic glycolysis system.

**Table 4:** Time Series Averages of Glucose, Lactate, and Pyruvate [54].

| Time (hours) | Lactate (mM) | Glucose (mM) | Pyruvate (mM) |
| --- | --- | --- | --- |
| 0 | 2.8 | 1.87 | 0.042 |
| 0.5 | 13.39 | 7.98 | 0.1014 |
| 1 | 21.78 | 10.49 | 0.1358 |
| 2 | 21.55 | 9.81 | 0.1360 |
| 3 | 21.38 | 9.62 | 0.1469 |
| 4 | 21.98 | 8.49 | 0.1362 |

To account for the distinct molecular dynamics and the increased proliferation rates specific to cancer cells, we considered slightly different values for ten parameters than those for a healthy cell. See Table 3 for the ten parameter values labeled as estimated. The different parameters showed that a cancer cell undergoes aerobic gycolysis in a more disorganized manner while the aerobic glycolysis process in a cone cell is more controlled. The different parameter values in cancer revealed less controlled lactate transport in and out of the cell, with significantly slower lactate transport contributing to ACoA, a faster pace of cell growth, a slightly higher glucose flux, lower ability to self-regulate glycolysis through citrate inhibition or less abundance of ATP, and less production of intermediate metabolites for ATP production by citrate. The differences in these mechanisms are defined by a much lower *k* value (0.45 versus 10) in the gating functions *h*([LACT]) and *f* ([LACT]), which illustrates back flow and not a complete on-off gating mechanism of lactate exchange between the extracellular and intracellular space with a significantly smaller *γ*, velocity of lactate transport into the cell for ACoA production; a higher fraction *q* of glucose converted to G3P for lipid synthesis and cell growth, which confirms the rapid cancer cell division and growth; a slightly larger range of glucose transport conversion factor λ, indicating more variability of glucose supplied from the RPE, including a higher demand for glucose; smaller citrate inhibition of glycolysis *β*, signifying less self-regulation or potentially less abundance of ATP; and smaller rate *ϕ* of converting citrate to ATP, illustrating a reduction in ATP. The lack of tight metabolic regulation in cancer was further shown by the two fold increase in external lactate L_*E*_, and the faster metabolic reaction of [CIT], given by the value of *V*_[CIT]_, and the larger *K*_[CIT]_ substrate concentration that gives half the maximal rate of *V*_[CIT]_.

The model simulations show a good fit with the data, with all three concentrations stabilizing to their steady states within a little over an hour, as shown in Figure 3. Our model assumes a constant external glucose flow from the RPE allowing for steady levels of [G] to be achieved while the experimental data comes from cultured cells leading to an eventual decay in [G]. Though there are many similarities in metabolism between cancer cells and photoreceptors in that both cell types exhibit the Warburg effect, retinal cell parameters differ. However, this qualitative match to data is a good proof of concept for our model, which can now be tuned to parameters specific for photoreceptors.

**Figure 3:**
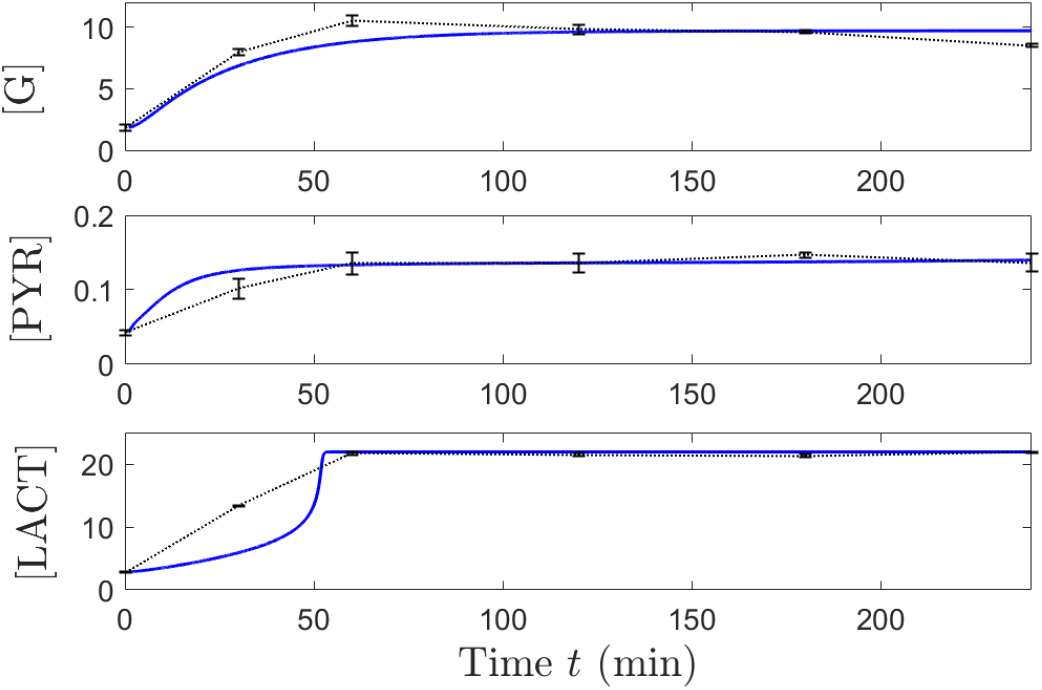
Fitting model predictions with data from cancer cells. In each panel, the black curve shows the average empirical values, with error bars describing variability (standard deviation) over a population of three measured cells [54]. The blue curve represents our predicted solution. The parameters used are as follows: λ = 0.0755, *ρ* = 0.05, *ξ* = 8, *δ* = 65, *α* = 0.2, *β* = 1, *q* = 0.38, *n* = 0.001, G_E_ = 13, *k* = 0.45, *γ* = 0.015, *ϕ* = 0.5, L_E_ = 22, *V*_[G]_ = 1.2, *K*_[G]_ = 19, *V*_[G3P]_ = 0.15, *K*_[G3P]_ = 0.143, *V*_[PYR]_ = 0.15, *K*_[PYR]_ = 1.7, *V*_[LACT]_ = 0.14, *K*_[LACT]_ = 0.125, *V*_[ACoA]_ = 0.15, *K*_[ACoA]_ = 0.02, *V*_[CIT]_ = 0.1, *K*_[CIT]_ = 0.5. Initial conditions for the simulation were chosen to agree with the average empirical ones (in mM): [G] = 1.87; [G3P] = 0.12; [PYR] = 0.042; [LACT] = 2.8; [ACoA] = 0.03; [CIT] = 0.02.

### 3.2 Bifurcation analysis and bistability ranges

As expected, the long-term dynamics of the system depend on its parameter values, and is altered by parameter perturbations. The model’s sensitivity to changes and uncertainty in its parameters, which define various key mechanisms on the cone metobolic system, are further analyzed in Section 3.3. Here, we observe the effects of perturbing specific key parameters, and discuss the crucial consequences of the number and position of steady states (which correspond to specific physiological states and may distinguish between viability or failure of the system).

We first analyzed the changes in dynamics in response to variations in the external glucose concentration, *G_E_*. Figure 4 shows the system’s equilibria and their evolution and phase transitions as *G_E_* is increased within the range of 0-13 mM. Each panel illustrates separately the projection of the same equilibrium curve along each of variable in the system, representing key metabolite concentrations. The figure suggests that a reduced extracellular glucose supply below 2.6 mM (i.e., *G_E_* < 2.6 mM) cannot successfully sustain the system and elevate internal glucose to a viable range. In this regime of *G_E_* < 2.6 mM, the only attainable long-term physiological state, represented by the only stable equilibirum reachable from any initial conditions, is a “low functioning” stable state, shown as a green solid curve in the figure, and characterized by [LACT] ~ 10 mM with all other metabolites concentrations close to zero. This represents a pathological state of the cone.

**Figure 4:**
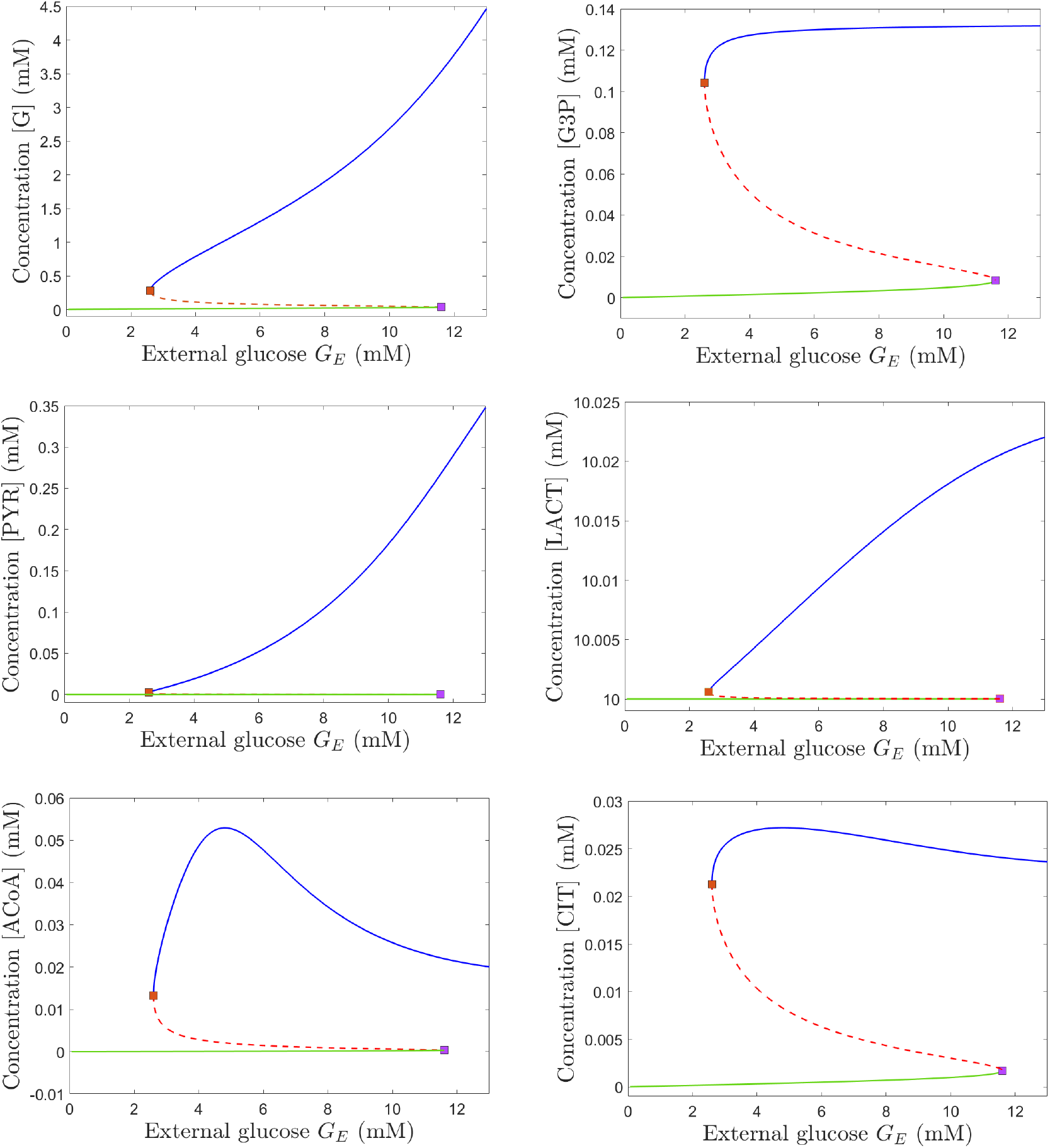
Equilibrium curves and bifurcations with respect to *G_E_*. As the level of external glucose is varied between *G_E_*=0-13 mM, the equilibria of the system are plotted, each panel representing a different component of the same equilibrium curves. There are two locally stable equilibrium branches shown as green and blue solid curves, and a saddle equilibrium, shown as a dotted red curve. The bistability window onsets with a saddle node bifurcation at *G_E_* ~ 2.6 mM (brown square marker), and closes with another saddle node bifurcation at *G_E_* ~ 11.6 mM (purple square marker). The other system parameters were held fixed as: λ = 0.0755, *n* = 0.001, *δ* = 65, *q* = 0.18, *β* = 1, *α*=0.2, *ρ* = 0.05, *φ* = 1, *ψ* = 8, *k* = 10, *γ* = 1, *L_E_* = 10; *V*_[G]_ = 1.2, *K*_[G]_ = 19, *V*_[G3P]_ = 0.15, *K*_[G3P]_ = 0.143, *V*_[PYR]_ = 0.15, *K*_[PYR]_ = 1.7, *V*_[LACT]_ = 0.14, *K*_[LACT]_ = 0.125, *V*_[ACoA]_ = 0.15, *K*_[ACoA]_ = 0.02, *V*_[CIT]_ = 0.03, *K*_[CIT]_ = 0.0054.

At *G_E_* ~ 2.6 mM, the system undergoes a saddle node bifurcation. If the external glucose level is raised past this phase transition value, the system enters a bistability regime, where a second, “viable” physiological steady state becomes available, with metabolite concentration levels in all components within a range for a healthy cone cell (illustrated in our panels as a blue solid branch of the equilibrium curve). Depending on the initial concentrations of the six metabolites in our model, the cone cell metabolism may go to the pathological or healthy state. The [G], [G3P], [PYR], [LACT], [ACoA], and [CIT] levels change in response to *G_E_* being further increased up to 13 mM. The internal glucose concentration [G] increases (up to ~ 4.5 mM), and so does the steady state level of [PYR], all the other components remain relatively unaltered, after a transient following up the birth of the second steady state. This shows the importance of external glucose and the components that alter it in driving the system via glucose and pyruvate metabolism.

The bistability window persists up to *G_E_* ~ 11.6 mM, allowing different initial conditions to converge to one of two locally attracting equilibria (the green and the blue curves, separated by the unstable saddle shown as a red dotted curve). Convergence of different initial states in different attraction basins to either of the two stable steady states is further illustrated in Figure 5. We show a [G]-[LACT] phase space slice, for a value of *G_E_* within the bistability range. For a more complete illustration, Figure 6 shows two potential evolutions of the system in the bistability regime (for *G_E_* = 10 mM). The left panel illustrates all components of the solution for a set of initial conditions in the basin of attraction of the green (“low functioning” or unhealthy) stable state, and the right panel for the blue (“high functioning” or healthy) steady state. The bistability window ends at *G_E_* ~ 11.6 mM, and henceforth the healthy equilibrium remains the only attainable state in the long run. The basin of attraction provides a range for the initial concentration levels of our six metabolites that will drive the system to either the pathological or healthy state depending on the parameter values. Investigating how varying the parameters leads to one of these two states provides potential mechanisms that can be altered as potential therapies for improving cone vitality and sight.

**Figure 5:**
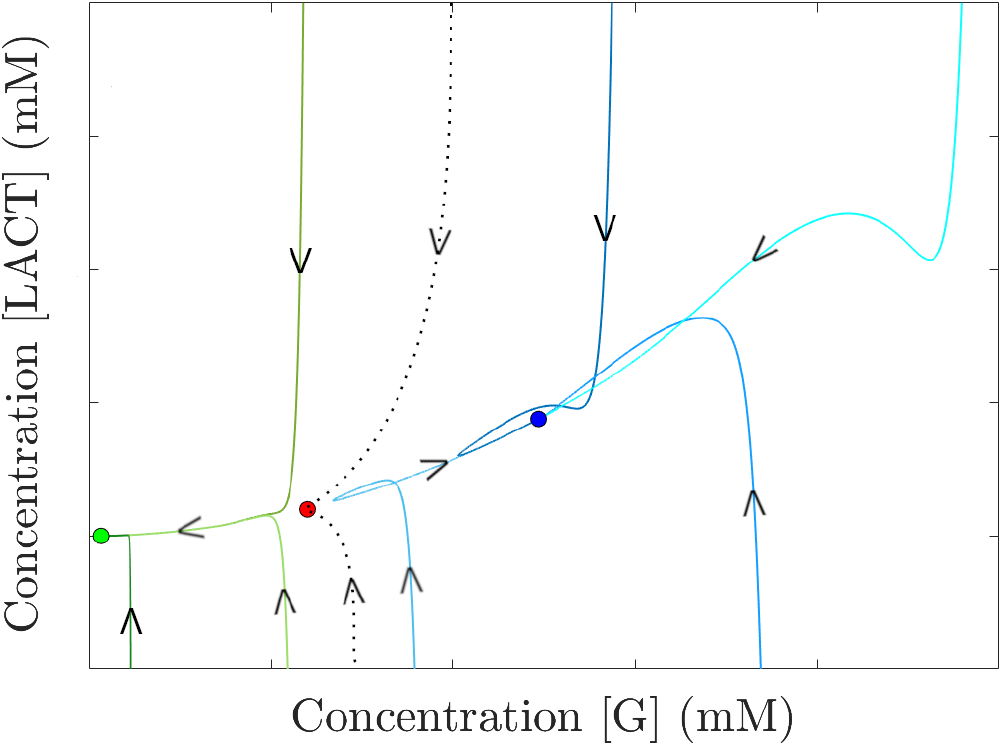
Schematic representation of coexistence of equilibria within the bistability window, shown in a phase-space two-dimensional slice [G]-[LACT]. The two stable equilibria are shown as a green and a blue dot. A third, saddle equilibrium is shown as a red dot. A few simulated trajectories converging to the green equilibrium are shown as green curves, and simulated trajectories which converge to the blue equilibrium are shown as blue curves. The stable manifold of the saddle was symbolically drawn as a dotted black curve. The fixed parameters are the same as in Figure 4. Figure 6 provides a more complete representation of all components for two representative solutions corresponding two different initial conditions; one converging to the green dot, and one converging to the blue dot, for *G_E_*=11.5 mM.

**Figure 6:**
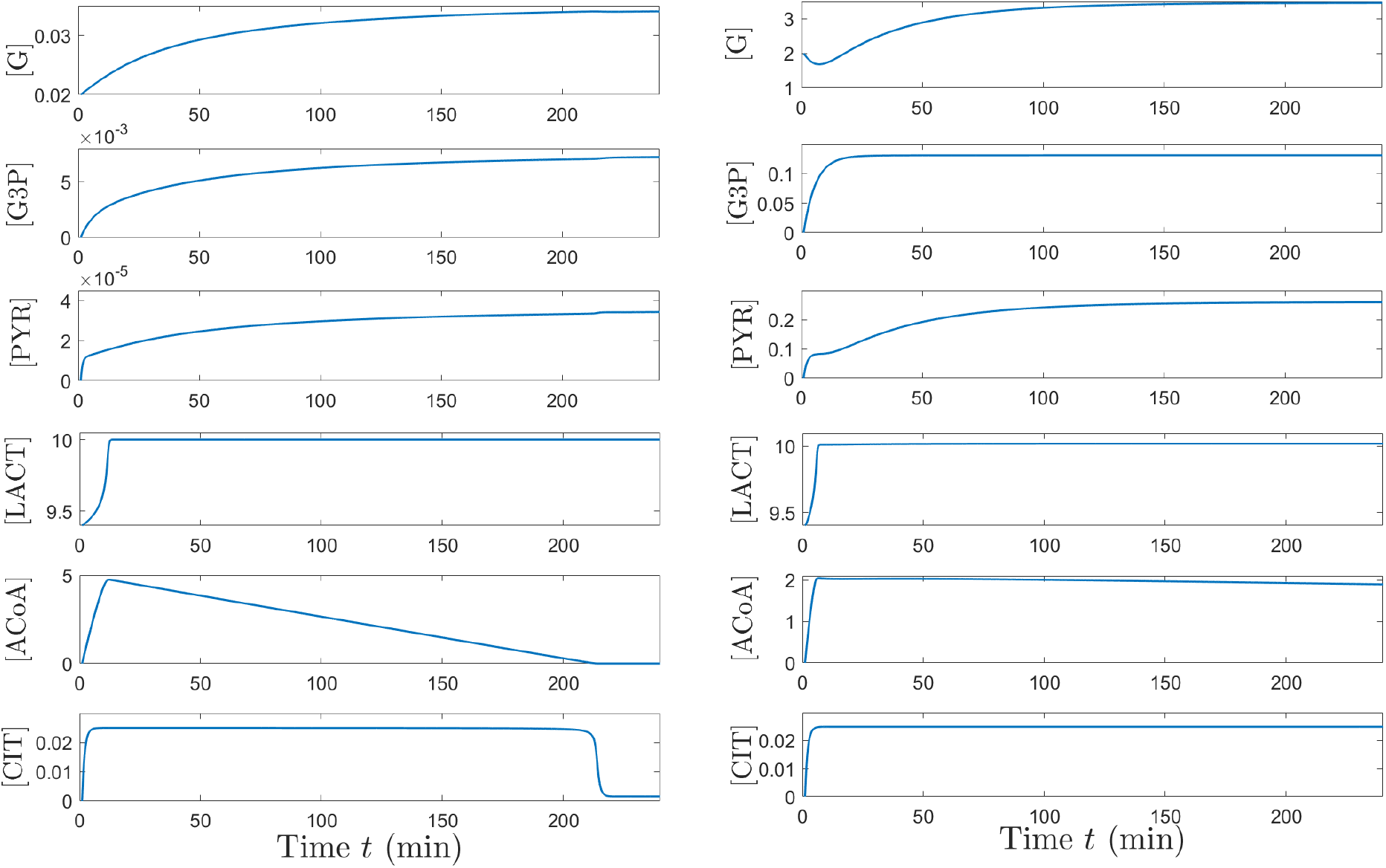
**Simulation of two solutions converging to the two different locally attracting equilibria**, for our system in the bistability regime (with external glucose concentration *G_E_*=11.5 mM and the rest of the parameters as in Figures 4) and 5. The left versus the right panels represent two trajectories which differ only in their initial glucose concentration level. The other initial concentration are [G3P]=[PYR]=[ACoA]=[CIT]=0, and [LACT]=9.4 mM. **Left.** Initial glucose level [G]=0.02 mM.The system converges to a non-viable steady state in which all concentrations are close to zero, except for lactate. **Right.** Initial glucose level [G]=2 mM. The system converges to a biologically viable/ healthy steady state as observed in empirical studies.

Tracking the behavior of the system in response to varying the transport conversion factor λ, or the rate of passive glucose transport *n*, leads to very similar bifurcation diagrams, bistability windows, and variable ranges. Thus, they are not further illustrated here. Instead, we focus on *α*, the rate of *β*-oxidation of ingested OS fatty acids created from G3P. While there is a bistability regime that lives between two saddle node points, the evolution of the system when varying *α* through these phase transitions is qualitatively different. We illustrate this behavior in Figure 7.

**Figure 7:**
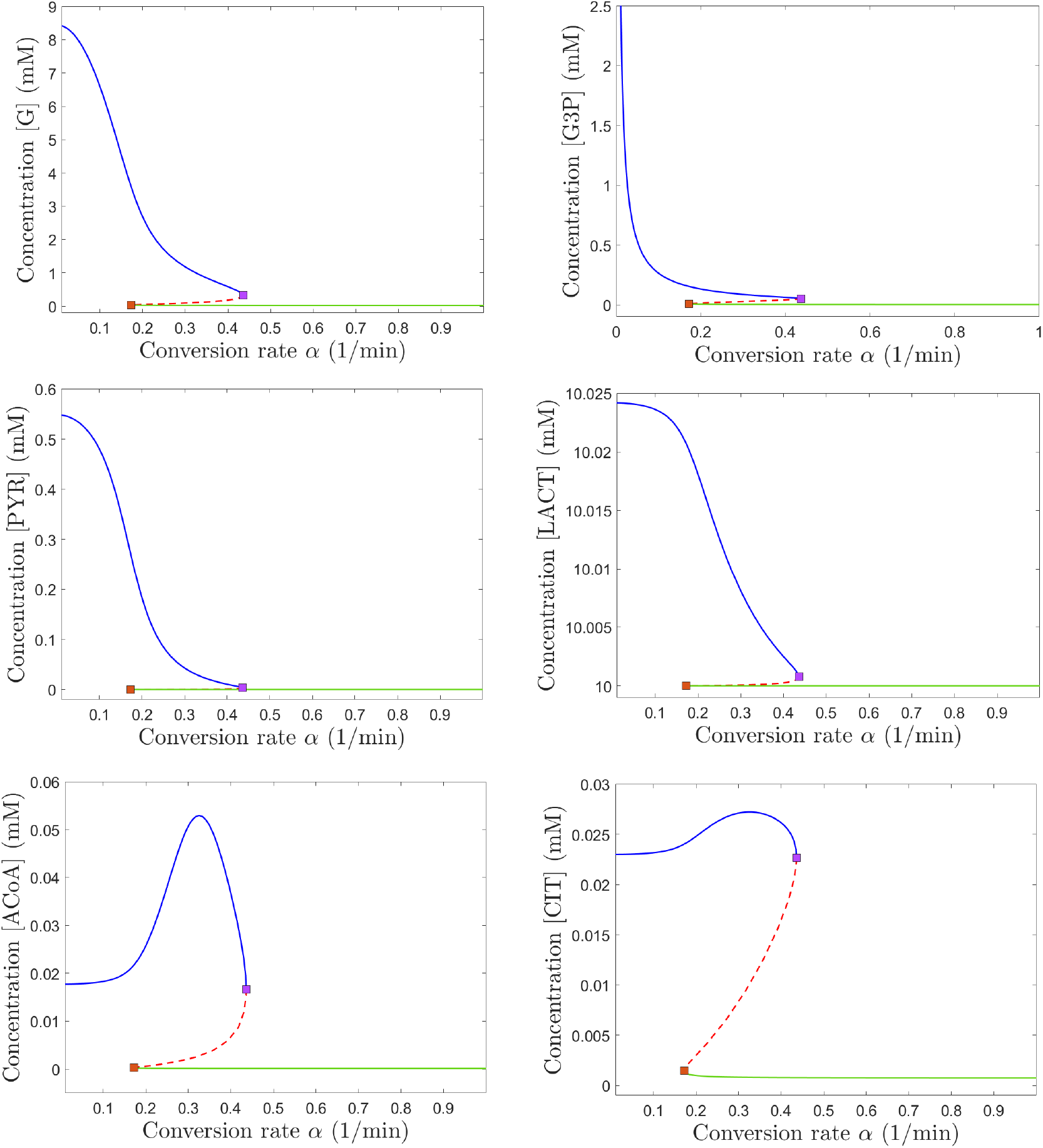
Equilibrium curves and bifurcations with respect to *α*. As the rate *α* is varied between 0 and 1 min^−1^, the equilibria of the system are plotted, each panel representing a different component of the same equilibrium curves. There are two locally stable equilibrium branches shown as green and blue solid curves, and a saddle equilibrium, shown as a dotted red curve. The bistability window onsets with a saddle node bifurcation at *α* ~ 0.17 min^−1^ (brown square marker), and closes with another saddle node bifurcation at *α* ~ 0.43 min^−1^ (purple square marker). *G_E_* was fixed to 10 mM. The other system parameters were held fixed as in Figure 4.

When the rate of *β*-oxidation of fatty acids (*α*) exceeds the bifurcation value 0.43 min^−1^, the system exhibits a unique locally stable equilibrium (solid green curve). This is a low functioning/unhealthy equilibrium, in the sense that all system components stabilize close to zero, except for [LACT] which stabilizes close to 10 mM, the external lactate value. Our analyses reveal that prolonged high rates of *β*-oxidation (beyond 0.43 min^−1^) which exist under extreme glucose starvation and scarce key metabolites will result in the pathological unhealthy state without any alternative for reprogramming the cone to a healthy state by altering certain processes or mechanisms. Further, our findings show that the gating mechanisms of lactate transport in the cone cell is a tightly controlled mechanism and thus always stabilizes close to the external lactate concentration.

Varying *α* above the bistability regime does not have significant impact on the long term outcome. However, when *α* is reduced past the saddle node bifurcation (purple marker), the system suddenly enters its bistability regime and gains access to a second, high functioning/ healthy steady state (blue solid curve). Rates of *β*-oxidation higher than 0.17 min^−1^ and lower than 0.43 min^−1^ provide the possibility of reprogramming the cone to a healthy state by altering certain processes and mechanisms. When *α* is decreased past the lower saddle node bifurcation at ~ 0.17 min^−1^, the green curve disappears in the collision with the unstable equilibrium, and the high functioning steady state becomes the only stable long term outcome. This result confirms that low rates of *β*-oxidation are aligned with a robust healthy metabolic state for the cone that can not be perturbed.

Since the blue curve represents the healthy viable outcome, and in fact the only stable outcome for small enough values of *α*, it is useful to track its progression in response to perturbations of the parameter. As *α* is progressively lowered, there is first an increase in all steady state components of the system. After an initial upward and then downward transient, the [ACoA] and [CIT] concentrations will settle to the same relatively low states ([ACoA] ~ 0.02 mM and [CIT] ~ 0.025 mM) as *α* approaches zero. The other steady state components will continue to increase as *α* approaches zero. While [G], [PYR], and [LACT] still settle to values in the biological range, [G3P] exhibits a blowup as *α* approaches zero. This is not at all surprising, since the [G3P] concentration is the compartment affected most directly by the shutdown of pathway **k** (i.e., by reducing to zero the *β*-oxidation of ingested OS fatty acids created from G3P). Under starvation or additional need of energy, *β*-oxidation of fatty acids becomes a key substrate to fuel ATP production in the TCA cycle. The bifurcation analysis for *α* shows that when initial [G], [PYR], and [G3P] levels are relatively low, *α* has the ability to change the fate of the cone cell and its metabolism. But *α* can only do this within a small range of values. This shows that this process of creating energy via intermediate substrates created from *β*-oxidation of fatty acids is mainly an auxiliary process and the main process by which the cell relies on intermediate metabolites and substrates.

### 3.3 Sensitivity Analysis

We use sensitivity analysis to determine which processes have the greatest impact on the intracellular concentrations tracked by the model. Sensitivity analysis includes the following general steps: i) vary the model parameters, ii) perform model simulations, iii) collect information on an output of interest (this can be the model output or another outcome), and iv) calculate sensitivity measures. There are local and global sensitivity analysis methods. In local methods, parameters are varied one at a time, and in global methods, all parameters are varied at the same time. Examples of global sensitivity analysis methods include Latin Hypercube Sampling/Partial Rank Correlation Coefficient (LHS/PRCC), the Sobol method, and Extended Fourier Amplitude Sensitivity Test (eFAST). Depending on the technique used, sensitivity measures are called coefficients or indices, and they indicate the impact of parameter changes on the output of interest [34].

In LHS/PRCC, the sensitivity measure is named the partial rank correlation coefficient (PRCC). This method can only be applied when parameter variations result in monotonic changes in the output [7, 34]. However, the advantages of LHS/PRCC compared to other global methods are simplicity and much lower computational demand. The magnitude of the PRCC values provides information about parameter influence on the outcome of interest. If the PRCC magnitude is greater than 0.4, the outcome of interest is considered sensitive to changes in the corresponding parameter [34]. The sign of a PRCC value shows if the corresponding parameter and the output are directly or inversely related. A positive coefficient indicates that the parameter and the output move in the same direction. A negative coefficient means they move in opposite directions, so as a parameter increases, the output decreases, and vice versa [7, 34]. In LHS/PRCC, parameters are varied simultaneously using Latin hypercube sampling (LHS). This involves assigning a probability distribution to each parameter, dividing the distribution into areas of equal probability and drawing at random and without replacement a value from every area [7, 34]. With LHS/PRCC, we can examine how a specific output is affected by an increase or decrease in a specific parameter, which can be useful for identifying the best parameters to target for control. Additionally, with LHS/PRCC we can explore how changes in initial conditions influence an outcome of interest [34].

The eFAST method can be conducted in the case when there are non-monotonic relationships between parameters (i.e., inputs) and a specific output of interest, but this approach is more computationally expensive than LHS/PRCC. In eFAST, the sensitivity measures are called sensitivity indices and they quantify the portion of variance in the outcome due to uncertainties in the parameters. There is a first order sensitivity index and total order sensitivity index. The first order index is a measure of how a parameter contributes to the output variance individually. The total order index shows the contribution a parameter makes to the output variance individually and in interaction with other parameters. The magnitude of sensitivity indices determines the importance of parameters [34, 45, 44]. In eFAST, parameters are varied at the same time using a sinusoidal search curve, where angular frequency is specified for each parameter. To compute the sensitivity indices for a given parameter, a high frequency is assigned to that parameter, while all other parameters are assigned a low frequency [34, 45, 44]. With eFAST, we can examine which parameter uncertainties have the largest impact on output variability [34]. Due to the intricacies and complexity of eFAST, initial conditions are rarely used as input factors. In the next section, we present the results of our sensitivity analysis using both the LHS/PRCC and eFAST methods.

### 3.4 Sensitivity Results

The results of our sensitivity analysis are summarized in Table 5, which correspond to the detailed results shown in Figures 8-10. For the normal photoreceptor model, parameters are varied over their corresponding ranges given in Table 3. For the case of cancer conditions, due to insufficient information regarding parameter ranges, we allowed for 30% variation around nominal values.

**Table 5:** PRCC and eFAST results on the model applied to a cancer cell and a normal cone cell for *t_f_* = 240 (minutes) with initial conditions [LACT]=9.4, [G3P]=[PYR]=[ACoA]=[CIT]=0 for cone cell and [G]=1.87, [G3P]=0.12, [PYR]=0.042, [LACT]=2.8, [ACoA]=0.03, and [CIT]=0.02 for cancer conditions. The sign of the PRCC is shown in parentheses with a threshold for significance being 0.4 and *p* < 0.01, and both first order and total order sensitivity indices considered in determining significance for eFAST. Both are presented in decreasing (magnitude) order of significance with bold font representing |PRCC| ≥ 0.7 and comparably large magnitude for the parameters using eFAST.

| Sensitivity index | Parameters with significant sensitivity index for stated conditions |  |  |
| --- | --- | --- | --- |
| (A) Uncertainty and sensitivity (US) analysis with [G] as outcome of interest over the different initial conditions |  |  |  |
|  | Normal with [G](0) = 0.02 | Normal with [G](0) = 2 | Cancer conditions |
| eFAST | $\alpha, G_E, K_{[G]}, V_{[G]}, V_{[PYR]}$ | $\alpha, G_E, K_{[G]}, V_{[G]}, V_{[PYR]}$ | $G_E, q, \alpha, \delta, \lambda, K_{[G]}, V_{[G]}$ |
| PRCC | $K_{[G3P]}, G_E$ | $\alpha(-), G_E, V_{[PYR]}(-), K_{[G]}(-), q, V_{[G]}(-), K_{[PYR]}$ | $G_E, \delta, \lambda, V_{[G]}, q, \alpha(-), K_{[G]}(-), V_{[G3P]}$ |
| (B) Uncertainty and sensitivity (US) analysis with [G3P] as outcome of interest over the different initial conditions |  |  |  |
|  | Normal with [G](0) = 0.02 | Normal with [G](0) = 2 | Cancer conditions |
| eFAST | $\alpha$ | $\alpha$ | $\alpha, V_{[G3P]}, q$ |
| PRCC | $\alpha(-), G_E, K_{[PYR]}$ | $\alpha(-), q, V_{[G3P]}$ | $\alpha(-), V_{[G3P]}, q$ |
| (C) Uncertainty and sensitivity (US) analysis with [PYR] as outcome of interest over the different initial conditions |  |  |  |
|  | Normal with [G](0) = 0.02 | Normal with [G](0) = 2 | Cancer conditions |
| eFAST | $\alpha, V_{[PYR]}, K_{[G]}, G_E, V_{[G]}$ | $\alpha, V_{[PYR]}$ | $V_{[LACT]}, V_{[PYR]}, q$ |
| PRCC | $G_E, K_{[PYR]}(-), K_{[G3P]}$ | $\alpha(-), V_{[PYR]}, K_{[G]}(-), q$ | $q(-), V_{[LACT]}(-), V_{[PYR]}, K_{[LACT]}$ |
| (D) Uncertainty and sensitivity (US) analysis with [LACT] as outcome of interest over the different initial conditions |  |  |  |
|  | Normal with [G](0) = 0.02 | Normal with [G](0) = 2 | Cancer conditions |
| eFAST | $L_E, \alpha, K_{[G]}$ | $L_E, \alpha, K_{[G]}$ | $L_E, k, V_{[PYR]}$ |
| PRCC | $[LACT]_0$ | $L_E$ | $L_E$ |
| (E) Uncertainty and sensitivity (US) analysis with [ACoA] as outcome of interest over the different initial conditions |  |  |  |
|  | Normal with [G](0) = 0.02 | Normal with [G](0) = 2 | Cancer conditions |
| eFAST | $L_E, \gamma, \alpha$ | $L_E, \alpha, \gamma$ | $L_E, k, q, \gamma, V_{[CIT]}$ |
| PRCC | $L_E, [LACT]_0(-), \gamma$ | $L_E, [LACT]_0(-)$ | $L_E, k, V_{[CIT]}(-), q, V_{[G3P]}$ |
| (F) Uncertainty and sensitivity (US) analysis with [CIT] as outcome of interest over the different initial conditions |  |  |  |
|  | Normal with [G](0) = 0.02 | Normal with [G](0) = 2 | Cancer conditions |
| eFAST | $L_E, \phi, V_{[CIT]}$ | $L_E, \phi, V_{[CIT]}$ | $\phi, L_E, V_{[CIT]}, k, V_{[G3P]}, q$ |
| PRCC | $\phi(-), V_{[CIT]}, L_E$ | $\phi(-), V_{[CIT]}, L_E$ | $\phi(-), L_E, V_{[CIT]}, q$ |

**Figure 8:**
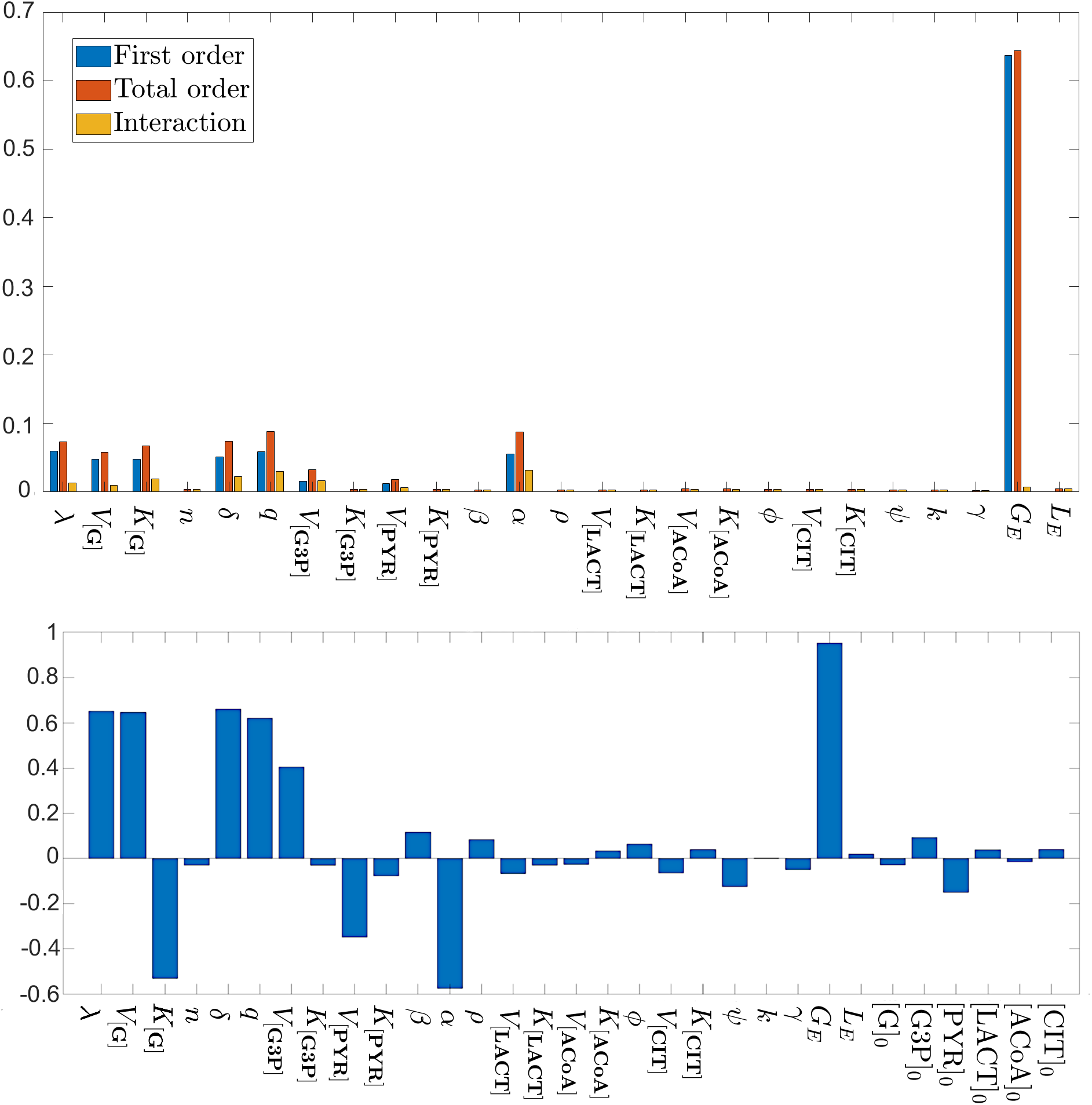
Sensitivity results for [G] using eFAST and LHS/PRCC for cancer conditions. These are the graphical results that are entered in the rightmost column of Table 5 for [G]. Both methods show that [G], the glucose level inside the cell, is most sensitive to changes in the parameter *G_E_*. Inspection of the top and bottom graphs shows comparable relative impact on [G] for the remaining parameters, with the exception of *V*_[G3P]_ which stands out as influential using PRCC but not eFAST.

#### Cancer conditions

Both the LHS/PRCC and the eFAST sensitivity analysis results for cancer conditions show that the glucose level inside the cell, [G], is most sensitive to changes in the parameter *G_E_*, the concentration of glucose outside the cell. The two methods also classify as important *q* (the fraction of glucose converted into G3P), *δ* (the increased uptake of glucose facilitated by hypoxia inducible factor 1 signaling, which up-regulates the expression of the glucose transporter GLUT1, for cancer cells [24]), *V*_[G]_ (the maximum transport rate of glucose), λ (the transport conversion factor), *α* (the rate of *β*-oxidation of ingested OS fatty acids [created from G3P]), and *K*_[G]_ (the substrate concentration giving half the maximal rate of *V*_[G]_). These parameters are involved in three key processes responsible for cell energy and growth; total glucose uptake (a catalyst in both), the utilization of G3P in *β*-oxidation (that results in *β*-HB which can be used as an oxidative substrate in the TCA cycle), and the production of G3P for lipid synthesis (which is essential for growth). The PRCC results also reveal that changes in the glucose concentration are inversely related with changes in the parameters *α* and *K*_[G]_; see Figure 8.

The PRCC and eFAST analyses for cancer conditions both reveal that the pyruvate concentration, [PYR], is sensitive to variation in the parameters which capture the maximum production rate of lactate and pyruvate inside the cone (*V*_[LACT]_ and *V*_[PYR]_, respectively) and the fraction of glucose converted into G3P (*q*); see Table 5. The negative PRCC values corresponding to the sensitivity of [PYR] to changes in *q* and *V*_[LACT]_ indicate that as the fraction of glucose diverted into G3P and the maximum production rate of lactate decrease, the pyruvate concentration, [PYR], increases. The PRCC approach also highlights how [PYR] is affected by *K*_[LACT]_, which measures the pyruvate concentration that gives half the maximal rate of *V*_[LACT]_. Changes in the chemical reaction of [LACT], in particular change in the mechanisms within as defined by parameters *V*_[LACT]_ and *K*_[LACT]_, affect the resulting [PYR] levels.

According to both sensitivity analysis methods, the lactate concentration level inside a cancer cell, [LACT], is significantly influenced by the concentration of lactate outside the cell, *L_E_*, both in its overall levels and its variability. The eFAST approach highlights two additional parameters that affect the variability of [LACT]. Uncertainty in the maximum production rate of pyruvate, *V*_[PYR]_, and the binding affinity level of the lactate transporter, *k*, will result in the variability of [LACT]. The sensitivity results for [G], [PYR], and [LACT] indicate that the initial biochemical reactions in the gycolysis pathway are sensitive to more mechanisms, as illustrated by the number of parameters in the corresponding cases in the third column of Table 5, than the reactions further downstream not including the reactions in the TCA and the Kennedy pathways.

#### Normal cone cell

Both the LHS/PRCC and eFAST methods show that for a cone cell, where model simulations are performed with initial condition for glucose of [G](0) = 0.02, the glucose concentration is sensitive to changes in the level of external glucose (*G_E_*). The results from the eFAST approach indicate that uncertainty in the rate of *β*-oxidation of ingested OS fatty acids (created from G3P) (*α*) has the more significant impact in the variability of [G]. For low initial concentration levels of glucose there are more mechanisms affecting [G] variability as indicated by the eFAST results. The LHS/PRCC and eFAST sensitivity results also have other differences. While PRCC highlights the G3P concentration that gives the half-maximum rate of *V*_[G3P]_ as having an impact on the intracellular glucose concentration, this parameter is not classified as important by eFAST. On the other hand, eFAST indicates that [G] is sensitive to variation in the parameters *K*_[G]_, *V*_[G]_, and *V*_[PYR]_.

The eFAST results using a higher initial condition for glucose of [G](0) = 2 show that the processes important for the intracellular glucose level are *β*-oxidation of OS fatty acids, external glucose, glucose uptake, and pyruvate production. In addition to indicating the impact of these factors, the PRCC method highlights the influence of converting glucose to G3P. For [G](0)=2, the same parameters impact [G] in the eFAST results. However, for PRCC the number of mechanisms affecting [G] increases so now changes in seven parameters (as opposed to two) affect [G]. These parameters are involved in glucose uptake, [PYR] biochemical reaction, glucose diversion to G3P, and *β*-oxidation. The negative sign of *α* in the PRCC analysis indicates that the glucose level in a cone cell decreases as the cell breaks down OS fatty acids at a higher rate to synthesize *β*-HB to be utilized as a substrate in the production of ACoA.

According to the eFAST results with [G](0) = 0.02, the pyruvate concentration, [PYR], in a cone cell is influenced by the oxidation of OS fatty acids (*α*), the amount of glucose outside the cell (*G_E_*), the half limiting value of the glucose transport rate (*K*_[G]_), the maximum rate of glucose transport (*V*_[G]_), and the maximum production rate of pyruvate (*V*_[PYR]_). In PRCC only three parameters affect [PYR]; *G_E_*, *K*_[PYR]_, and *K*_[G3P]_. The substrate that gives the half-maximal rate of *V*_[PYR]_, defined by *K*_[PYR]_, inversely affects [PYR]. An increase in *K*_[PYR]_ will increase the amount of substrate required for [PYR] to reach its saturation level.

The eFAST and PRCC results with [*G*](0) = 2 differ from those with lower initial condition for glucose: eFAST no longer classifies external glucose and glucose uptake as influential, and PRCC shows a whole new set of parameters as being important. In addition, there are less mechanisms (defined by the model’s parameters) affecting [PYR] in the eFAST results as compared with the LHS/PRCC results for [*G*](0) = 2. Both sensitivity methods highlight [PYR] as being affected by changes and uncertainties in the parameters that describe *β*-oxidation and maximum production rate of [PYR], *α* and *V*_[PYR]_, respectively. The PRCC results indicate an inverse relationship between variation in *α* and *K*_[G]_ and changes in [PYR]. PRCC also identifies the process of diverting glucose to G3P for production of OS, which are rich in fatty acids, as having a strong effect. See Figure 9 for the case with initial condition [G](0) = 2.

**Figure 9:**
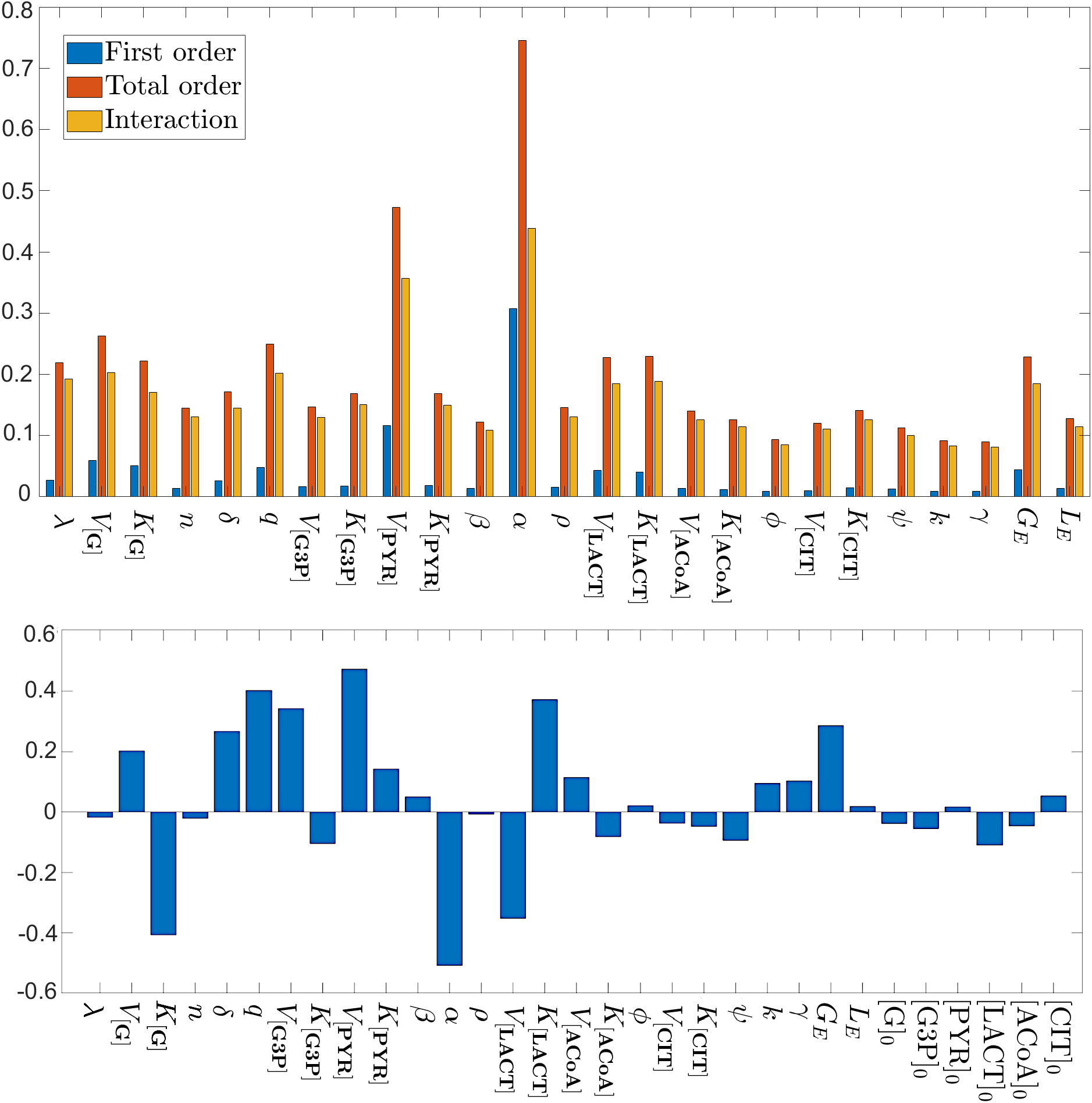
Sensitivity results for [PYR] using eFAST and LHS/PRCC using normal photoreceptor conditions with [G](0) = 2. These are the graphical results that are entered in the relevant column of Table 5 for [PYR]. The graphs illustrate agreement in the importance of *α* and *V*_[PYR]_.

When the intracellular lactate level, [LACT], is the outcome of interest, both sensitivity analysis methods show as important the extracellular lactate level (*L_E_*) when [G](0)=2. The eFAST results also classify the processes of *β*-oxidation of OS fatty acids from lipids produced by G3P (*α*) and the half-limiting value of glucose transport (*K*_[G]_) as having an impact on [LACT]. The eFAST results for [LACT] were the same for [G](0)=0.02 and [G](0)=2. The PRCC results show that only the lactate initial condition has an impact on the concentration of lactate inside the cell when the initial internal glucose concentration is low; see Figure 10 for the case of with initial condition [G](0) = 0.02. The relatively small number of parameters that affect [LACT] levels and variability indicates the strong pull of these mechanisms (or parameters) to try to bring the external and internal lactate levels to a balance.

**Figure 10:**
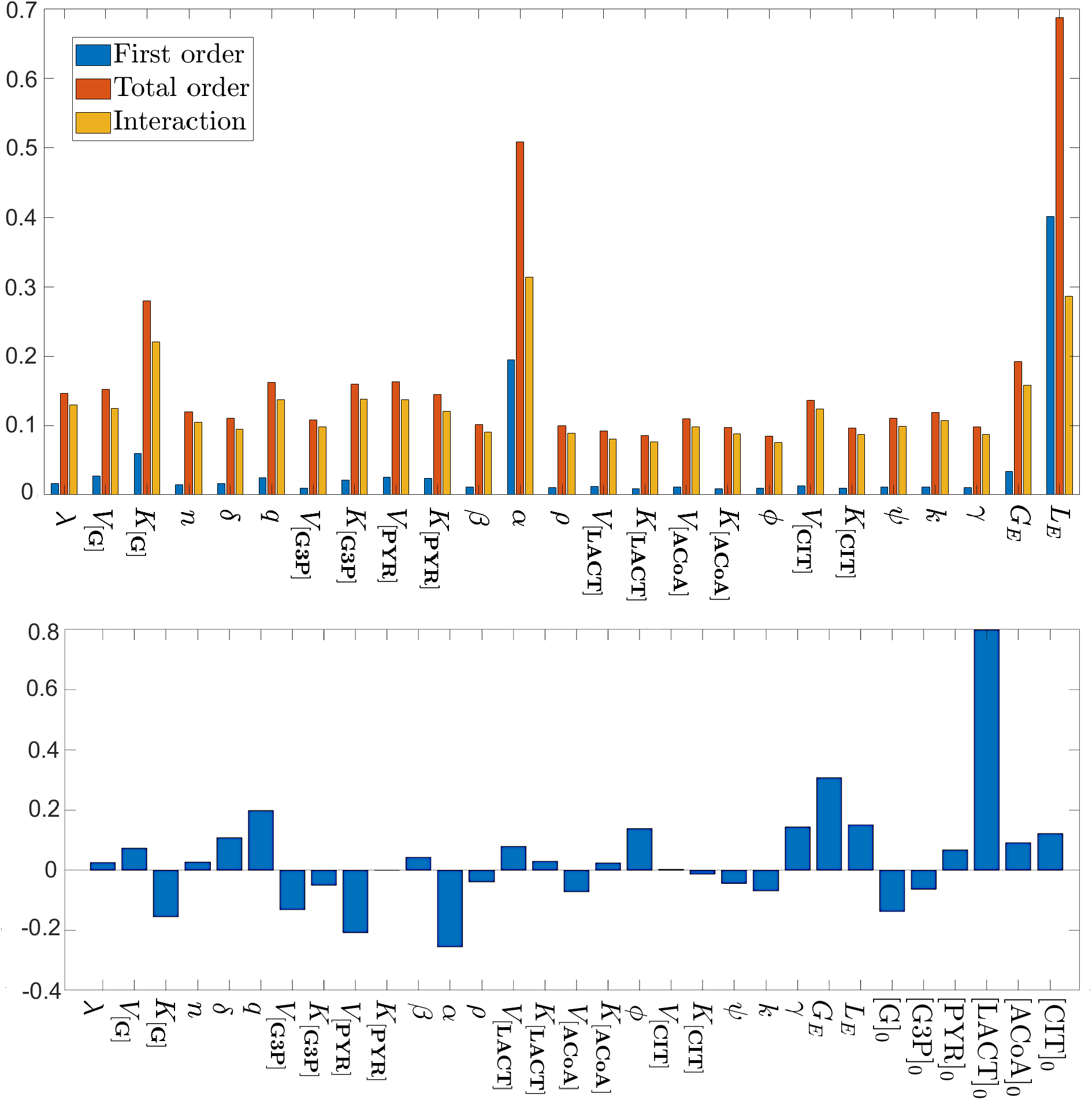
Sensitivity results for [LACT] using eFAST and LHS/PRCC using normal photoreceptor conditions with [G](0) = 0.02. These are the graphical results that are entered in the relevant column of Table 5 for [LACT]. While eFAST classified *L_E_* and *α* as most influential, PRCC determined that the initial lactate concentration is most important.; see the text for further discussion of these results.

For both [G](0)=0.02 and [G](0)=2, PRCC and eFAST indicate that *β*-oxidation of OS fatty acids from lipids produced by G3P (*α*) is the most important process for the level and variability of [G3P]. In addition, PRCC shows that external glucose (*G_E_*) and the half-limiting value of pyruvate production (*K*_[PYR]_) have an impact on the [G3P] level for [G](0)=0.02, while at the higher initial condition for glucose, important factors are the conversion of glucose to G3P (*q*) and the half-limiting value of G3P production (*K*_[G3P]_).

The eFAST results show that uncertainty in the same parameters influences the variability of [AcoA] at both [G](0)=0.02 and [G](0)=2. These parameters are *L_E_* (external lactate level), *α* (*β*-oxidation of OS fatty acids), and *γ* (maximum velocity of lactate transport contributing to AcoA). PRCC also highlights *γ* as important but only for [G](0)=0.02. With both initial conditions for glucose, the PRCC results indicate that the external lactate level and the initial internal lactate concentration have a strong impact on the level of ACoA.

The sensitivity results of [CIT] for [G](0)=0.02 and [G](0)=2 are the same for both methods. PRCC and eFAST indicate that [CIT] is impacted by external lactate (*L_E_*), rate of CIT conversion to ATP (*ϕ*), and the maximum production rate of CIT (*V*_[CIT]_). The relative impact of these mechanisms differs within each method. Uncertainties in external lactate affect the variability of [CIT] the most in eFAST, with the rate of CIT conversion to ATP having the second largest impact. PRCC reveals that *ϕ* affects [CIT] the most, with *V*_[CIT]_ having the second largest impact. An increase in the rate of CIT conversion to ATP reduces the concentration of CIT, while an increase in the maximum production rate of CIT elevates [CIT].

Interestingly, across both the normal photoreceptor model and cancer conditions, changes in *α* affect [G] and [G3P] but only [PYR] in the cone cell. In an analogous manner, changes in external lactate, *L_E_*, affect [LACT], [ACoA], and [CIT] across photoreceptor and cancer conditions (with one exception).

## 4 Discussion

### 4.1 Specific comments on the model

In this work, we developed and explored a mathematical model for the dynamics in the metabolic pathways of a healthy photoreceptor cell. We validated our model structure by comparing its predictions for concentrations of glucose, lactate, and pyruvate to data collected in cancer cells [54], which are metabolically similar to photoreceptors. In addition to developing the model structure, we also identified parameter values and ranges through a comprehensive literature search. When possible, we used values specific to photoreceptor cells and, if no measurements existed, selected another cell type as a proxy.

We applied two different global sensitivity analysis methods (LHS/PRCC and eFAST) and found the sensitive parameters resulting from each. PRCC reveals how the output of a model is affected if a parameter is changed, whereas variance-based methods such as eFAST measure which parameter uncertainty has the largest impact on output variability [34]. Using these two sensitivity analysis approaches in unison, we obtained a comprehensive view for which processes reflected in the equations (via the parameters) have the greatest impact on the metabolic system.

This sensitivity analysis, for the case of a photoreceptor cell, revealed that external glucose (reflected by *G_E_*) and *β*-oxidation of fatty acids from OS (generated by G3P lipid synthesis) for ACoA production (defined by *α*) significantly affect the concentrations of glucose, G3P, and pyruvate at steady state (corresponding to time equal to 240 minutes). The importance of external glucose indicates that the effective metabolism of photoreceptors relies on sufficient availability of glucose, their primary fuel resource. The influence of *β*-oxidation of fatty acids, which links glycolysis occurring in the cytosol to oxidative phosphorylation in the mitochondria, suggests that photoreceptor metabolism is modulated by this feedback mechanism. The PRCC results also indicate that at a low initial level of intracellular glucose, the pyruvate concentration is most sensitive to changes in external glucose, while if greater amount of intracellular glucose is present initially, the pyruvate concentration is most sensitive to changes in the rate of *β*-oxidation.

Our sensitivity analyses reveal that ACoA, CIT, and intracellular lactate are impacted to the greatest extent by external lactate (*L_E_*). Furthermore, they are the only metabolites sensitive to external lactate. This suggests that external lactate is an important mechanism affecting oxidative phosphorylation, while it does not seem to have a strong influence on the glycolysis pathway, where external glucose has a crucial role.

We used bifurcation techniques to study the dependence of the system’s behavior on the parameters, in particular on *G_E_* and *α*, identified as key parameters by the sensitivity analysis. We found that the system undergoes two saddle node bifurcations with respect to these parameters revealing bistability over a range of parameter values. This is heartening, as a properly designed analysis should reveal bifurcation parameters to be sensitive [34]. Bistability allowed us to investigate the mechanisms, defined by the parameters, that can be altered to bring a cone to healthy conditions from the pathological state. We were also able to determine key ranges for *G_E_* and *α* as well as initial metabolite levels that will lead to one state versus another with the aid of bifurcation curves and basins of attraction.

Our analysis found that the system behaves monotonically (broadly speaking) as the external glucose concentration is increased. This is not surprising when considering the molecular coupling: as more glucose is made available to the cell, the internal glucose concentrations are expected to increase, driving in turn (via pathways **c** and **d**) higher concentrations of G3P and PYR, respectively, and further (via pathway **g**) a higher concentration of LACT. The levels of [LACT] eventually increase above the external concentration *L_E_*, preventing additional production of [ACoA], and subsequently of [CIT].

Our bifurcation analysis also reveals that a very low external source of glucose (less than 2.6 mM) cannot drive the cell to function in a healthy regime (since in the long term, all metabolites will be depleted without an adequate source of glucose to maintain cone metabolism, except for the *β*-HB and external lactate used as substrates for ACoA and fuel for ATP production.

Increasing *G_E_* to an adequate level pushes the system into the bistability window, with two potential, and very different outcomes. This opens up the possibility for the cell to function in a healthy long-term regime (with concentrations which have been observed empirically within the healthy functional range for the eye). Such alternative prognosis is available based on the cone cell’s current state or ability to change the current metabolite levels in the cell. If all the molecular pools of the cell have become extremely low, the cell can no longer be rescued by increasing the extracellular glucose. Figure 4 illustrates how an external concentration *G_E_* =11.5 mM or lower cannot resuscitate a cell with already depleted molecular pools [G3P]=[PYR]=[ACoA]=[CIT]=0, [LACT] close to the external concentration of 10 mM, and [G]=0.02 mM. The same external concentration of *G_E_* =11.5 mM can bring a cell to a healthy regime provided the initial glucose is as high as [G]=2 mM. Such a surge of glucose input may have, however, other physiological effects on the system, not captured by our model, and should not be necessarily viewed as a cure-all strategy.

Our sensitivity and bifurcation analyses support the expectation from the model diagram (Figure 2) that the system’s prognosis also depends quite crucially on the rate *α* of *β*-oxidation of ingested OS fatty acids. The metabolites directly affected by even small changes in *α* are [G], [G3P], [PYR], [LACT], and [ACoA], but these perturbations propagate via the tight coupling of the system, affecting the long-term concentrations of all its components.

Our bifurcation analysis in Section 3.2 shows the global effects of having an overly glucose-starved system corresponding to an extremely large rate of *β*-oxidation. Overall, too high of a *β*-oxidation rate leads to a complete system shutdown. An extremely low *β*-oxidation rate leads to a dangerously high accumulation of G3P in the cell, as the system under-utilizes lipids to be converted to OS (whose fatty acids will eventually be utilized to generate *β*-HB).

The bifurcation analysis for *α* between ~ 0.17-0.43 mM, shows that the bistability regime, hence the optimal functioning of the system, is contingent on its initial state. If the current state of the cell is close to healthy, further tuning its *β*-oxidation rate (e.g., via medication or therapies) can optimize its function. However, if the cell’s current state is very poor (e.g., based on a history of functioning under pathological parameter values), the behavior cannot be rescued even by a substantial adjustment in the *β*-oxidation rate, and the cell’s function will remain poor. These effects support our sensitivity analysis, which showed all metabolites of the system (with the exception CIT) to be sensitive to changes or perturbation of *α*.

It has been established that the RPE serves as the principal pathway for the exchange of metabolites (in particular, glucose and lactate) between the choroidal blood supply and the retina [14]. The fact that external glucose (*G_E_*) and the rate of *β*-oxidation of fatty acids (*α*) are highlighted as important by our sensitivity and bifurcation analyses, as well as external lactate (*L_E_*) in the sensitivity analyses, points to the critical role the RPE plays in photoreceptor metabolism. As these processes link the metabolism of photoreceptors with the metabolism of the RPE, our findings indicate that the normal function of photoreceptors relies heavily on their interaction with the RPE. This aligns with the physiological understating that photoreceptors and the RPE have a reciprocal resource relation and operate as a functional unit: the RPE provides photoreceptors with a source of metabolism via glucose, and photoreceptors provide a source of metabolism for the RPE via lactate.

External lactate is key for maintaining a balanced reciprocal resource relation between the RPE and photoreceptors, on which cone nutrition and vitality depend. In addition to glucose supplied by the RPE, photoreceptors can also consume lactate, produced by other retinal cells for oxidative metabolism. For example, Müeller cells are known to secrete lactate which can be used as fuel in photoreceptors [49]. The high impact of external lactate seen in the sensitivity analysis also points to the importance of this mechanism for photoreceptor metabolism.

### 4.2 Limitations and future work

We have identified three limitations in our model. (1) Our model considers a single healthy photoreceptor cell, whereas in reality there are multiple photoreceptors of different types and other cell types such as RPE and Müeller cells, forming a “metabolic ecosystem.” Future work will address this complexity. (2) Our model would be improved if time series data existed for concentration levels of the metabolites, represented by state variables, in the retina. We identified time series concentrations of glucose, lacate, and pyruvate for cancer cells, which are metabolically similar to photoreceptors, but we would expect parameters to differ. Time series data would also allow us to better estimate parameters to which our model outputs are sensitive. (3) Some parameter ranges were not available from the literature, so we were forced to use a different tissue type as a proxy. Because both LHS/PRCC and eFAST depend on starting with biologically relevant ranges for each parameter and sampling within that range, this is a concern. If photoreceptor-specific parameter ranges become available, this model could be updated and improved.

The RPE is a layer of cells which provides glucose to photoreceptors, and these cells are also metabolically active. External glucose is very important, and though it is a parameter in our single-cell model, in future work it will depend on the dynamics of other cell types. The RPE serves as the main pathway for the exchange of critical metabolites (specifically, glucose and lactate) between the choroidal blood supply and the retina [14]. Metabolites that can be used as substitutes for photoreceptor energy production, during glucose deprivation, are also mediated by the RPE. Müeller cells are known to secrete lactate which can be used as fuel in photoreceptors [49]. A future step in this work will be investigating the interaction of the RPE, Müeller cells, and photoreceptors along with the “metabolic ecosystem” they create.

## 5 Conclusions

We developed and analyzed a mathematical model for the dynamics in the metabolic pathways of a healthy photoreceptor cell. Using two different methods for sensitivity analysis, we identified the parameters and potential mechanisms that are driving system output levels and variability which are particularly relevant to photoreceptor health. The behavior of the model for different values of the highly sensitive parameters was explored, and we demonstrated that certain sets of parameters exhibit phase transitions and bistable behavior where healthy and pathological states both exist.

Our work confirms the necessity for the external glucose, *β*-oxidation, and external lactate concentrations, which are key feedback mechanisms connecting the RPE and photoreceptors, to sustain the cell. The role of *β*-oxidation of fatty acids which fuel oxidative phosphorylation under glucose- and lactate-depleted conditions, is validated. A low rate of *β*-oxidation corresponded with the healthy cone metabolite concentrations in our simulations and bifurcation analysis. Our results also show the modulating effect of the lactate differential (internal versus external) in bringing the system to steady state; the bigger the difference, the longer the system takes to achieve steady state. Additionally, our parameter estimation results demonstrate the importance of rerouting glucose and other intermediate metabolites to produce glycerol 3-phosphate (G3P), to increasing lipid synthesis (a precursor to fatty acid production) to support their cone cell high growth rate. A number of parameters are found to be significant; however, the rate of *β*-oxidation of ingested outer segment is shown to consistently play an important role in the concentration of glucose, G3P, and pyruvate, whereas the extracellular lactate level is shown to consistently play an important role in the concentration of lactate and acetyl coenzyme A.

These mechanisms can be posed to the biology community for future experiments or for potential therapeutic targets. The ability of these mechanisms to affect key metabolites’ variability and levels (as revealed in our analyses) signifies the importance of inter-dependent and inter-connected feedback processes modulated by and affecting both the RPE’s and cone’s metabolism. The modeling and analysis in this work provide the foundation for a more biologically complex model that metabolically couples different cell types as found in the retina.

## 6 Acknowledgements

This work was initiated during the Association for Women in Mathematics collaborative workshop Women Advancing Mathematical Biology hosted by the Institute of Pure and Applied Mathematics at University of California, Los Angeles in June 2019. Funding for the workshop was provided by IPAM, NSF ADVANCE “Career Advancement for Women Through Research-Focused Networks”(NSF-HRD 1500481). We are grateful to Nancy Philp for introducing us to the importance of lactate consumption in photoreceptor metabolism and for fruitful discussions related to this work. We would like to thank the two anonymous reviewers for their thoughtful comments that made made the manuscript clearer and stronger.

## A Simple Model

In this section, System (1)-(6) is simplified to gain more insight into the cone glucose metabolic pathways. The model is simplified as below:

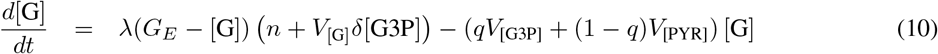

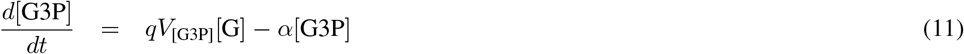

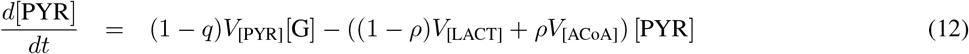

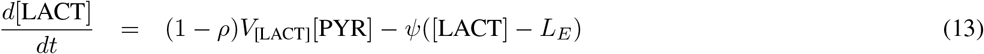

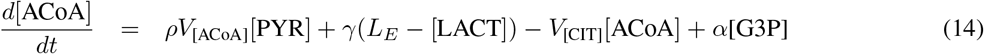

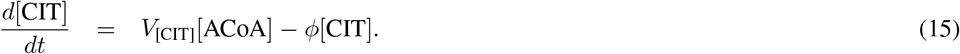

We were able to prove existence and uniqueness of the model solution in System (10)-(15). Furthermore, it was proved that the system evolves to a unique equilibrium point under healthy conditions. However, the simple model does not capture the complete qualitative behaviours of the full model.

### A.1 Model Analysis

Note that System (10)-(15) is well-posed and that all solutions remain within the state space, [G] ≥ 0, [G3P] ≥ 0, [PYR] ≥ 0, [LACT] ≥ 0, [ACoA] ≥ 0, [CIT] ≥ 0, since the right-hand side functions of System (10)-(15) are continuously differentiable [48]. The analysis of Model (10)-(15) is done by finding the equilibria and their corresponding stability properties. Setting the right-hand sides of the equations (10)-(15) equal to zero yields the following biological meaningful equilibrium point denoted by *E*([G*], [G3P*], [PYR*], [LACT*], [ACoA*], [CIT*]), and defined as follows

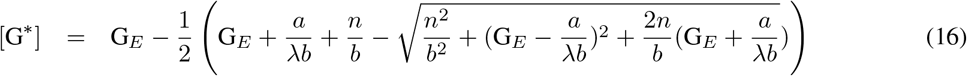

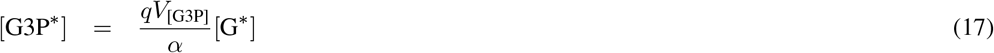

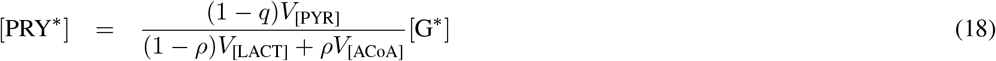

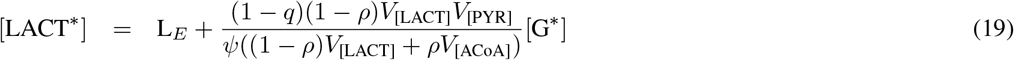

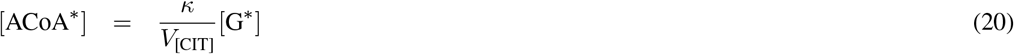

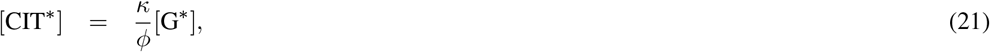

where *a* = *qV*_[G3P]_ + (1 − *q*)*V*_[PYR]_, *b* = *qδV*_[G]_*V*_[G3P]_*/α* and

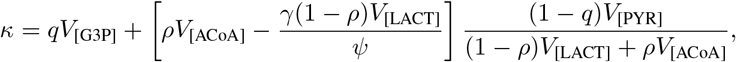

with *κ* ≥ 0.

#### Theorem 1.

*The equilibrium E exists and it is locally stable.*

*Proof.* From the parameter modeling assumptions, it is easy to prove that [G*] > 0. Therefore, Equations (17)-(19) are all positive and Equations (20)-(21) are non-negative if and only if *κ* ≥ 0. Hence, *E* is a biologically feasible equilibrium, since all the elements of *E* are non-negative for all parameter values of the model. Next, the Jacobian matrix corresponding to *E* is given by the following lower triangular block matrix:

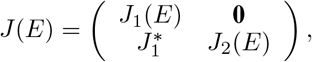

where 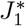 is a non-zero matrix and

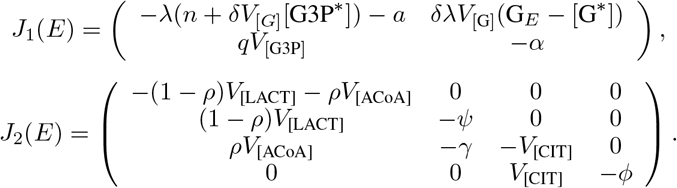

Therefore, the eigenvalues of *J*(*E*) are determined from the eigenvalues of *J*_1_(*E*) and *J*_2_(*E*). Since *J*_2_(*E*) is a lower triangular matrix, its eigenvalues are given by its diagonal elements, which are all negative by the parameter modeling assumptions. From Routh-Hurwitz criteria, *n* = 2, the eigenvalues of *J*_1_(*E*) are negative or have negative real part if and only if *det*(*J*_1_(*E*)) > 0 and *tr*(*J*_1_(*E*)) < 0. From the model assumptions follow that *tr*(*J*_1_(*E*)) = −λ(*n* + *δV*_[*G*]_[G3P*]) − *a* − *α* < 0 and

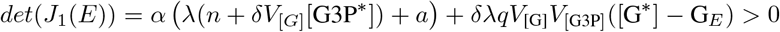

if an only if

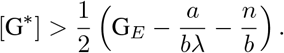

From Equation (16)

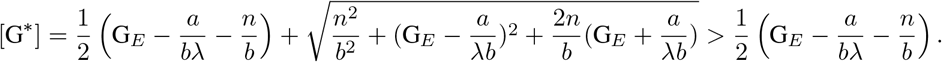

Therefore, all the eigenvalues of *J*(*E*) are negative and hence *E* is a locally stable node.

Therefore in a long term glucose metabolic dynamic behaviour within a single cone, all the substrate variables achieve steady-state values, which depend linearly on the steady-state glucose concentration value, [G*], Equation (17)-(21). Furthermore, the concentration of glucose inside of the cell is always less than the outside concentration while the lactate concentration inside is more than the outside concentration, i.e.,

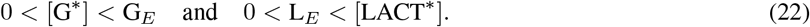

Note that [G*] = 0 if and only if *α* = 0 or *n* = 0 and 0 < *G_E_* < *a/bλ*. Therefore *α* and *n* are important parameters for the survival of the cell. Another important parameter is *κ*, since when *κ* = 0 the [ACoA*] = 0, and [CIT*] = 0 which also leads to a pathological metabolic steady-state outcome.

Figure 11 shows the molecular evolution of System (10)-(15), where the variables evolve to their steady-state values in about 3.3 hours (200 min).

**Figure 11:**
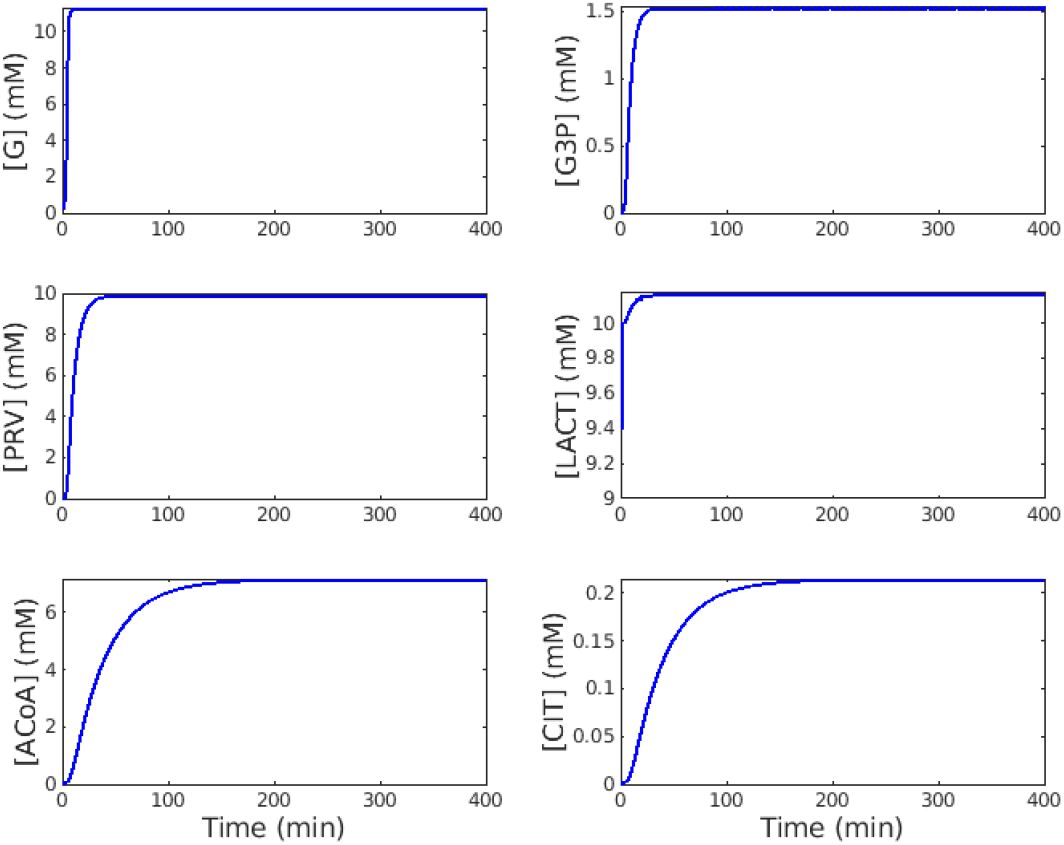
Molecular evolution of energy demand in a single cone cell. The parameter values are set to their baseline value for normal photoreceptor model with initial conditions: [G] = 0.2, [LACT] = 9.4 and the rest of the other initial conditions are set to be equal to zero.

## B Acronym Glossary

| Acronym Definitions |  |
| --- | --- |
| Acronym | Meaning |
| ACoA | acetyl coenzyme A |
| ATP | adenosine triphosphate |
| BSG-1 | basigin-1 |
| CIT | citrate |
| DHAP | dihydroxyacetone phosphate |
| eFAST | Extended Fourier Amplitude Sensitivity Test |
| G | glucose |
| GAP | glyceraldehyde-3-phosphate |
| G3P | glycerol-3-phosphate |
| G6P | glycerol-6-phosphate |
| GLUT1 | glucosetransporter 1 |
| LACT | lactate |
| LDH | lactate dehydrogenase |
| LDHA | lactate dehydrogenase A |
| LDHB | lactate dehydrogenase B |
| LHS | Latin Hypercube Sampling |
| MCT | monocarboxylate transport proteins |
| NAD | nicotinamide adenine dinucleotide |
| OS | outer segment |
| OXPHOS | oxidative phosphorylation |
| PDC | pyruvate dehydrogenase complex |
| PFK | phosphofructokinase |
| PRCC | Partial Rank Correlation Coefficient |
| PYR | pyruvate |
| RdCVF | rod-derived cone viability factor |
| RdCVFL | rod-derived cone viability factor long form |
| RPE | retinal pigmented epithelium |
| TCA | tricarboxylic acid |
| VEGF | vascular endothelial growth factor |

## Footnotes

1 There are approximately 20 rods per each cone in the human retina (and 25 to 1 in mice retina). G3P leads to the production of lipids which result in new photoreceptor OS. Thus we take [G3P] as a proxy for rods with the appropriate scaling factor incorporated into *δ*, the scaling factor for contribution of RdCVF by rods.

2 Since we are not considering the RPE, we will utilize *α*[G3P] as a proxy for the metabolite *β*-hydroxybutyrate produced by the PRE and utilized by the photoreceptor’s TCA.

3 By the inverse relation of the functions *f* ([LACT]) and *h*([LACT]), the parameters *k* and L_*E*_ have the analogous meaning with respect to each function.

4 The authors generously shared their data used to generate their Figure 1B for three cells for each experiment.

